# Subthalamic-Cortical Network Reorganization during Parkinson’s Tremor

**DOI:** 10.1101/2021.03.26.437170

**Authors:** Peter M. Lauro, Shane Lee, Umer Akbar, Wael F. Asaad

## Abstract

Tremor, a common and often primary symptom of Parkinson’s disease, has been modeled with distinct onset and maintenance dynamics. To identify the neurophysiologic correlates of each state, we acquired intraoperative cortical and subthalamic nucleus recordings from ten patients performing a naturalistic visual-motor task. From this task we isolated short epochs of tremor onset and sustained tremor. Comparing these epochs, we found that the subthalamic nucleus was central to tremor onset, as it drove both motor cortical activity and tremor output. Once tremor became sustained, control of tremor shifted to cortex. At the same time, changes in directed functional connectivity across sensorimotor cortex further distinguished the sustained tremor state.

**SIGNIFICANCE STATEMENT:** Tremor is a common symptom of Parkinson’s disease (PD). While tremor pathophysiology is thought to involve both basal ganglia and cerebello-thalamic-cortical circuits, it is unknown how these structures functionally interact to produce tremor. In this manuscript, we analyzed intracranial recordings from the subthalamic nucleus and sensorimotor cortex in patients with PD undergoing deep brain stimulation (DBS) surgery. Using an intraoperative task, we examined tremor in two separate dynamic contexts: when tremor first emerged, and when tremor was sustained. We believe that these findings reconcile several models of Parkinson’s tremor, while describing the short-timescale dynamics of subcortical-cortical interactions during tremor for the first time. These findings may describe a framework for developing proactive and responsive neurostimulation models for specifically treating tremor.

## INTRODUCTION

Tremor, a cardinal symptom of Parkinson’s disease (PD), typically manifests as a 4–6 Hz oscillatory movement of the distal limbs during rest or sustained posture (Lance et al., 1963). While often the presenting motor symptom of PD, tremor (and its response to dopamine replacement therapy) is highly variable across patients (Koller, 1984; Zach et al., 2015; Koller, 1986; Dirkx et al., 2017; Dirkx et al., 2019). PD tremor neurophysiology has been described by the “dimmer switch” model where an “on-off” mechanism is separable from a magnitude controller (Helmich et al., 2012). Specifically, functional MRI (fMRI) BOLD activity from basal ganglia nuclei such as the globus pallidus *pars interna* (GPi) correlates with the presence or absence of tremor, whereas immediate tremor amplitude better correlates with BOLD signal from structures in cerebello-thalamo-cortical circuits such as motor cortex (Helmich et al., 2011;Helmich, 2018). The GPi, and the monosynaptically-connected subthalamic nucleus (STN) (Albin et al.,1989), are common therapeutic targets for deep brain stimulation (DBS). Indeed, DBS in each nucleus is equally effective in reducing tremor (Wong et al., 2020). However, the precise role of the STN and its interactions with cortex in these tremor dynamics is unknown.

Low-frequency (4-8 Hz) oscillatory bursting has been observed in both in the STN and GPi in MPTP primate models of PD (Bergman et al., 1994; Raz et al., 2000). This bursting, although present in the absence of tremor, becomes highly synchronized with tremor once it emerges. STN recordings from patients with PD have similarly revealed *θ*/tremor-frequency (3-8 Hz) activity that is coherent with electromyography (EMG) recordings of tremulous limbs (Levy et al., 2000; Reck et al., 2009; Reck et al., 2010). Accordingly, STN tremor frequency oscillations (along with higher frequency oscillations) have been used to predict clinical measures of tremor (Hirschmann et al., 2016; Telkes et al., 2018;Asch et al., 2020). Further, studies applying STN DBS at tremor frequencies entrained tremor to the phase of the stimulation, consistent with a direct modulatory role of STN on tremor (Cagnan et al.,2014).

At the same time, tremor reorganizes cortical activity. Magnetoencephalography (MEG) studies of patients with PD identified a broad cortical tremor network comprising “intrinsic” (ventrolateral anterior thalamus (VLa), premotor and motor cortex) and “extrinsic” (cerebellum, ventrolateral intermedius (VIM), somatosensory cortex) loops hypothesized to initialize and stabilize tremor respectively (Volkmann et al., 1996; Timmermann et al., 2003). This cortico-cortical synchronization at single and double tremor frequencies extends to STN local field potential (LFP) and EMG recordings as well (Hirschmann et al., 2013). Meanwhile, intraoperative studies combining electrocorticography (ECoG) and STN LFP recordings found decreases in *α* (8–13 Hz) and *β* (13–30 Hz) coherence during tremor (Qasim et al., 2016). Despite this broad cortico-cortical synchronization at tremor frequencies, it remains unclear whether these neurophysiological changes are specific to tremor onset or maintenance. In addition, although STN and sensorimotor cortex become coherent during tremor, the manner in which tremor-related activity is coordinated across structures, and how different networks of activity may reflect the different stages of tremor production and maintenance, are unknown.

Thus, in order to understand whether there are indeed distinct neurophysiological mechanisms of tremor initiation and maintenance, and to better understand what neurophysiological interactions characterize these states, we recorded local field potential activity from the STN along with ECoG from sensorimotor cortices while subjects with PD engaged in a task that elicited initiation and persistence of tremor. Specifically, we tested whether the STN (like the GPi) drove tremor specifically during onset, while cortical structures drove sustained tremor.

## MATERIALS AND METHODS

### Participants

All patients undergoing routine, awake placement of deep brain stimulating electrodes for intractable, idiopathic PD between November 2015 and September 2017 were invited to participate in this study **(Table 1)**. Patients with PD were selected and offered the surgery by a multi-disciplinary team based solely upon clinical criteria, and the choice of the target (STN vs. GPi) was made according to each patient’s particular circumstance (disease manifestations, cognitive status and goals) (Akbar and Asaad,2017). In this report, we focused on ten patients (9M, 1F) undergoing STN DBS with intraoperative ECoG recordings. Patients were off all anti-Parkinsonian medications for at least 12 hours in advance of the surgical procedure (UPDRS Part III: 48.2 ± 15.6). Four patients were considered tremor-dominant, and six patients had average tremor UPDRS III scores > 2 in their right hand (Jankovic et al., 1990). Approximately age-matched controls (3M, 11F; often patients’ partners) also participated in this study (*n* = 14 subjects); patients were aged 55.6–78.5 years (65.2 ± 7.4), and controls were aged 48.3–79.2 years (62.4 ± 10.0) at the time of testing (Mann-Whitney U-test, *p* > 0.05). Controls were required simply to be free of any diagnosed or suspected movement disorder and to have no physical limitation preventing them from seeing the display or manipulating the joystick. There was a strong male-bias in the patient population (9M, 1F) and a female preponderance in the control population (3M, 11F), reflecting weaker overall biases in the prevalence of PD and the clinical utilization of DBS therapy (Accolla et al., 2007;Hariz et al., 2011; Rumalla et al., 2018). All subjects were right-handed. Patients and other subjects agreeing to participate in this study signed informed consent, and experimental procedures were undertaken in accordance with an approved Rhode Island Hospital human research protocol (Lifespan IRB protocol #263157) and the Declaration of Helsinki.

### Surgical Procedure

Microelectrode recordings (MER) from the region of the STN of awake patients are routinely obtained in order to map the target area and guide DBS electrode implantation. A single dose of short-acting sedative medication (typically propofol) was administered before the start of each procedure, at least 60–90 minutes prior to MER. The initial trajectory was determined on high-resolution (typically 3T) magnetic resonance images (MRI) co-registered with CT images demonstrating previously-implanted skull-anchor fiducial markers (version 3.0, FHC Inc., Bowdoin, ME, USA). Localization of the target relied upon a combination of direct and indirect targeting, utilizing the visualized STN as well as standard stereotactic coordinates relative to the anterior and posterior commissures. Appropriate trajectories to the target were then selected to avoid critical structures and to maximize the length of intersection with the STN. A 3-D printed stereotactic platform (STarFix micro-targeting system, FHC Inc.) was then created such that it could be affixed to these anchors, providing a precise trajectory to each target (Konrad et al.,2011). Microdrives were attached to the platform and then loaded with microelectrodes. Recordings were typically conducted along the anterior, center, and posterior trajectories (with respect to the initial MRI-determined trajectory) separated by 2 mm, corresponding to the axis of highest anatomical uncertainty based upon the limited visualization of the STN on MRI. Bilateral electrocorticography (ECoG) strips were placed posteriorly along sensorimotor cortices through the same burr hole used for MER insertion for temporary recordings. MER began about 10-12 mm above the MRI-estimated target, which was chosen to lie near the inferior margin of the STN, about 2/3 of the distance laterally from its medial border. The STN was identified electrophysiologically as a hyperactive region typically first encountered about 3-6 mm above estimated target (Gross et al., 2006). At variable intervals, when at least one electrode was judged to be within the STN, electrode movement was paused in order to assess neural activity and determine somatotopic correspondence, as per routine clinical practice. At these times, if patients were willing and able, additional recordings were obtained in conjunction with patient performance of the visual-motor task.

### Neurophysiological Signals and Analysis

Microelectrode signals were recorded using “NeuroProbe” tungsten electrodes (Alpha Omega, Nazareth, Israel). ECoG signals were acquired using Ad-Tech 8-contact subdural strips with 10 mm contact-to-contact spacing (Ad-Tech Medical, Racine, WI). All signals were acquired at 22-44 kHz and synchronized using Neuro Omega data acquisition systems (Alpha Omega). Microelectrode impedances were typically 400-700 kΩ while ECoG contact impedances were typically 10-30 kΩ. Patients performed up to 4 sessions of the task, with microelectrodes positioned at different depths for each session. As microelectrodes were not independently positionable, some signals may have necessarily been acquired outside of the STN. All recorded signals were nevertheless considered and analyzed.

Neural data were analyzed using the “numpy/scipy” Python 3 environment (Harris et al., 2020; Virtanen et al., 2020) (https://numpy.org/, https://www.scipy.org/). Offline, ECoG contacts were re-referenced to a common median reference within a strip (Liu et al., 2015). All resulting signals were bandpass filtered between 2-600 Hz, and notch filtered at 60 Hz and its harmonics. Timeseries were Z-scored and artifacts above 4 standard deviations were removed. These resulting timeseries were then downsampled to 1 kHz. Timeseries were bandpass filtered using a Morlet wavelet convolution (wave number 7) at 1 Hz intervals, covering 3–400 Hz. The instantaneous power and phase at each frequency was then acquired by the Hilbert transform. To analyze broad frequency bands, we grouped frequencies as: *θ*: 3–8 Hz, *α*: 8–12 Hz, *β_low_*: 12–20 Hz, *β_high_*: 20–30 Hz, *γ_low_*: 30–60 Hz, *γ_mid_*: 60–100 Hz, *γ_high_*: 100–200 Hz, and *hfo*: 200–400 Hz. For interregional analyses (phase-locking value, phase slope index, and granger prediction) we focused on frequencies up to 100 Hz; spectral or timeseries data were subsequently downsampled to 250 Hz.

### Anatomical Reconstruction of Recording Sites

Patients underwent pre-, intra- and post-operative imaging per routine clinical care. Preoperatively, stereotactic protocol magnetic resonance (MR) images were obtained (Siemens Vario 3.0 T scanner) that included T1- and T2-weighted sequences (T1: MPRAGE sequence; TR: 2530 ms, TE: 2.85 ms, matrix size: 512 × 512, voxels: 0.5 × 0.5 mm^2^ in-plane resolution, 224 sagittal slices, 1 mm slice thickness; T2: SPACE sequence, TR: 3200 ms, TE: 409 ms, matrix size: 512 × 512, voxels: 0.5 × 0.5 mm^2^ in-plane resolution, 224 sagittal slices, 1 mm slice thickness). Pre-, intra-, and post-operative (in some cases) computed tomography (CT) scans were also acquired (Extra-Op CT: GE Lightspeed VCT Scanner; Tube voltage: 120 kV, Tube current: 186 mA, data acquisition diameter: 320 mm, reconstruction diameter: 250 mm, matrix size: 512 × 512 voxels, 0.488 × 0.488 mm^2^ in-plane resolution, 267 axial slices, 0.625 mm slice thickness; Intra-Op CT: Mobius Airo scanner, Tube voltage: 120 kV, Tube current: 240 mA, data acquisition diameter: 1331 mm, reconstruction diameter: 337 mm, matrix size: 512 × 512 voxels, 0.658 × 0.658 mm^2^ in-plane resolution, 182 axial slices, 1 mm slice thickness). Postoperative MR images (Seimens Aera 1.5 T scanner, T1: MPRAGE sequence, TR: 2300 ms, TE: 4.3 ms, matrix size: 256 × 256 voxels, 1.0 × 1.0 mm^2^ in-plane resolution, 183 axial slices, 1 mm slice thickness, specific absorption rate < 0.1 W/g) were typically obtained 1–2 days after the operation to confirm proper final electrode location.

To reconstruct recording locations, MR and CT images were co-registered using the FHC Waypoint Planner software. The raw DICOM images and the linear transform matrices were exported and applied to reconstructed image volumes using the AFNI command “3dAllineate,” bringing them into a common coordinate space (Cox, 1996; Li et al., 2016). Microelectrode depths were calculated by combining intraoperative recording depth information with electrode reconstructions obtained from postoperative images using methods described previously (Lauro et al., 2015; Lauro et al., 2018). To determine the anatomical distribution of microelectrode recording sites across patients, preoperative T1-weighted MR images were registered to a T1-weighted MNI reference volume (MNI152_T1_2009c) using the AFNI command “3dQwarp”(Fonov et al., 2009). The resulting patient-specific transformation was then applied to recording site coordinates. MNI-warped recording coordinates were then assessed for proximity to structures such as the STN as delineated on the MNI PD25 atlas (Xiao et al., 2012; Xiao et al., 2015;Xiao et al., 2017). ECoG contacts were segmented from intraoperative CT volumes using the same DBStar processing as microelectrodes. Contacts were then projected onto individual cortical surface reconstructions generated from preoperative T1 volumes (Dale et al., 1999; Fischl et al., 2002; Saad and Reynolds, 2012; Trotta et al., 2018). Individual cortical surface reconstructions were co-registered to a standard Desikan-Destrieux surface parcellation (Argall et al., 2006; Desikan et al., 2006; Destrieux et al., 2010). Contacts were labeled and grouped as “premotor cortex,”“motor cortex,”“somatosensory cortex,” or “parietal cortex” if they contained the following anatomical parcellation labels:

- Premotor cortex/PMC : ctx_lh_G_frontjsup, ctx_lh_G_front_middle
- Motor cortex/MC : ctx_lh_G_precentral
- Somatosensory cortex/SC : ctx_lh_G_postcentral
- Posterior Parietal cortex/PPC : ctx_lh_G_parietal_sup, ctx_lh_G_pariet_inf-Supramar

If a contact had more than one label (8/80 contacts), they were removed from further analysis.

### Experimental Design

We employed a visual-motor target tracking task to estimate the degree of motor dysfunction in a continuous fashion. Specifically, while patients with PD reclined on the operating bed in a “lawn-chair” position, a joystick was positioned within their dominant hand, and a boom-mounted display was positioned within their direct line-of-sight at a distance of ~1 meter. The task was implemented in MonkeyLogic (Asaad and Eskandar, 2008a; Asaad and Eskandar, 2008b; Asaad et al., 2013) and required subjects to follow a green target circle that moved smoothly around the screen by manipulating the joystick with the goal of keeping the white cursor within the circle **(Figure 1A)**. The target circle followed one of several possible paths (invisible to the subject), with each trial lasting 10-30 seconds. Each session consisted of up to 36 trials (~13 minutes of tracking data), and subjects performed 1-4 sessions during the operation. Control subjects performed this task in an extra-operative setting.

**Figure 1.**
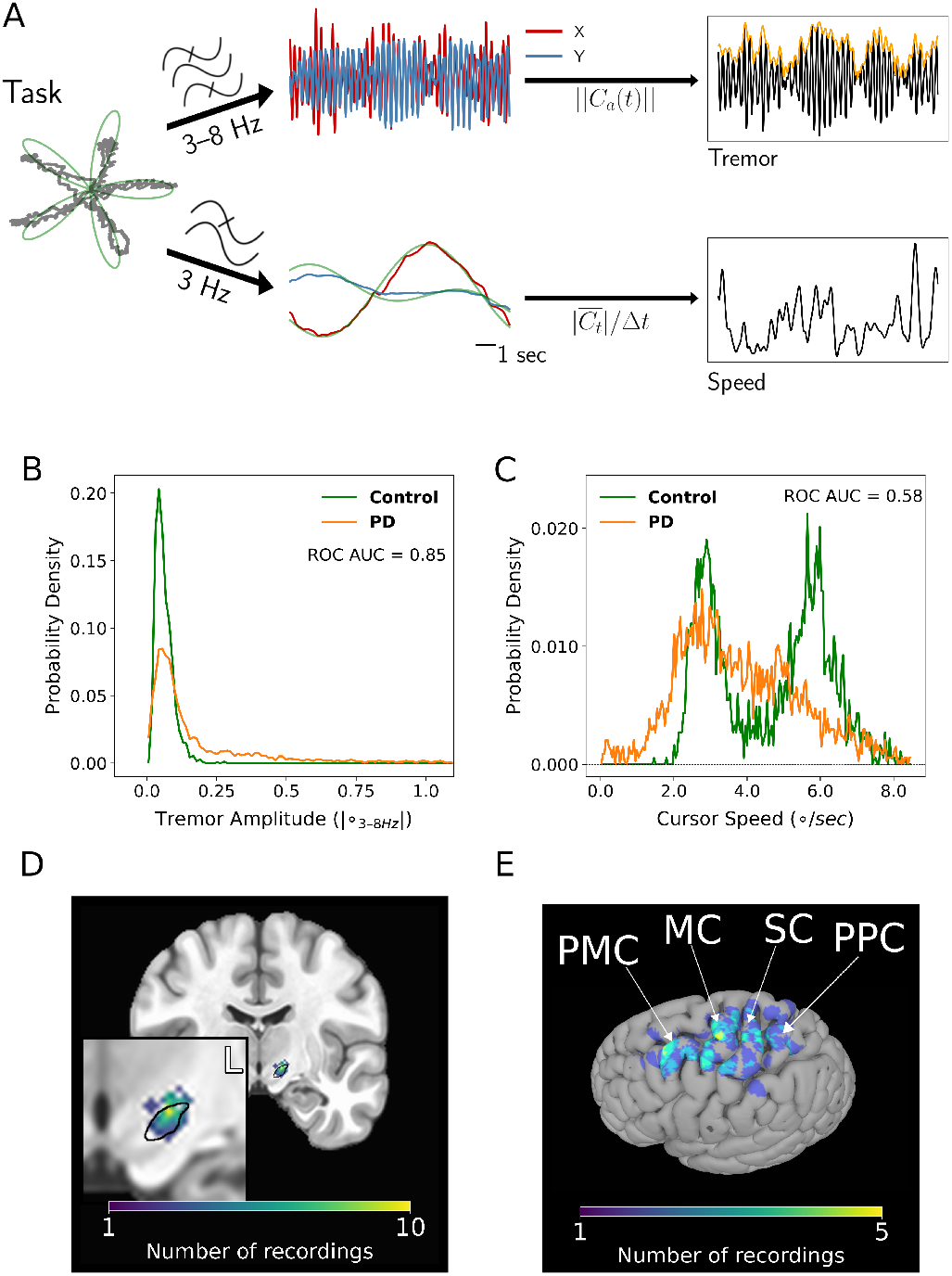
Tremor and movement speed calculated from the intraoperative visual-motor task. ***A***, Left - Schematic of task target (green) and joystick (gray) traces from a single trial. Center-top - Bandpass filtered X and Y joystick traces from the task trial. Center-bottom - Lowpass filtered X and Y joystick traces from the task trial. Right-top - One-dimensional projection of bandpass filtered traces (black), with tremor amplitude measured from the envelope (orange). Right-bottom - Cursor speed measured from lowpass filtered traces (black). ***B***, Distribution of 4 second tremor amplitude epochs for control subject and PD patient populations. ° - degrees of visual angle. Vertical dashed line indicates ROC-derived cutoff value between control and PD populations. While there is overlap on the left side of the distribution (patients with PD can exhibit control-like performance), the PD distribution is highly skewed on the right side of the distribution, allowing a large range of tremor expression. ROC AUC - Receiver operator characteristic area under the curve. ***C***, Distribution of 4 second speed epochs for control subject and PD patient populations. The bimodality of the control distribution corresponded to the pre-programmed speed of the onscreen target. Despite this, note that the PD distribution is shifted towards lower speed values. ***D***, Coronal view of microelectrode recording density on an MNI reference volume. The inset panel displays a close-up view of the subthalamic nucleus (outlined in black). L - left. ***E***, Recording density of ECoG contacts on an MNI reference surface. PMC - premotor cortex; MC - motor cortex; SC - somatosensory cortex; PPC - parietal cortex.

### Speed Quantification

To calculate movement speed, x- and y-joystick traces were 3 Hz low-pass filtered, and the euclidean change of cursor position was calculated over time. To standardize movement speed within patients, movement speed values within a session were min-max normalized into a measure of “slowness,” where 0=highest speed and 1=lowest speed.

### Tremor Amplitude Quantification

To calculate tremor, x- and y-joystick traces were 3-8 Hz bandpass filtered, and a one-dimensional linear projection of the filtered traces was calculated. Tremor amplitude and phase were calculated using the Hilbert transform of the resulting one-dimensional timeseries.

### Tremor Epoch Design

To standardize tremor amplitude across patients, tremor amplitude values from controls and patients were averaged into 4 second contiguous, non-overlapping epochs. The resulting average and standard deviation of the control tremor amplitude distribution were used to Z-transform control subject and PD patient tremor amplitude epochs **(Figure 1B)**. To determine a cutoff to optimally differentiate control and PD population tremor data, receiver operator characteristic (ROC) tests were performed between supra-cutoff population data for cutoff values ranging from −2 (the lowest observed in both populations) and 10. The maximum area-under-curve (AUC) value was observed for Z=3 (ROC AUC = 0.85), which was used for subsequent analyses.

To analyze neural activity associated with different tremor dynamics, 4 second tremor epochs were defined as following:

- “No Tremor” epochs were characterized by tremor staying below a 3 s.d. threshold for 4 seconds.
- “Tremor Onset” epochs were characterized by tremor exceeding a 3 s.d. threshold for 2 seconds, with tremor in the preceding 2 seconds being sub-threshold.
- “Sustained Tremor” epochs were characterized by tremor staying above a 3 s.d. threshold for 4 seconds.

All epochs were non-overlapping in time.

### Tremor Frequency Calculation and UPDRS Correlation

To calculate each patient’s dominant tremor frequency (i.e. the frequency with the largest amplitude), a distribution of tremor amplitude was created by aggregating each patient’s tremor amplitude epochs. In parallel, a frequency distribution was created by calculating the dominant tremor frequency within each epoch. A patient-specific dominant tremor frequency was then calculated as the frequency containing the highest aggregate tremor amplitude.

Correlations between task-derived tremor and UPDRS were conducted with sub-scores pertaining to the upper extremity relevant to the patient’s task performance (Rest Tremor, Postural Tremor, Finger Taps, Hand Opening/Closing, Rapid Alternating Movements (RAM), Rigidity). Each patient UPDRS sub-score was Spearman correlated with the median of each patient’s tremor amplitude distribution, and was assessed for significance using a bootstrap null distribution where tremor medians were randomly shuffled with respect to UPDRS sub-scores.

### Tremor/Speed-Spectral Power Correlation

To determine if spectral power across frequencies correlated with changes in tremor amplitude or slowness, Spearman correlations were calculated between 4 second epochs of averaged tremor/slowness and spectral magnitude of narrowbands with 1 Hz bandwidth. Correlations were calculated within entire task sessions. To determine whether spectral-tremor correlations were consistently positive or negative across all sessions, *ρ*-value distributions were tested for asymmetry about zero using Wilcoxon tests (Wilcoxon p-values corrected by the Benjamini-Hochberg procedure at *q* = 0.05).

### Tremor Epoch Spectral Power Modulation

To determine if spectral power at each structure differed by tremor epoch type, spectral power across frequencies were compared using the Kruskal-Wallis test. If spectral power in a frequency band was found to significantly differ across epoch types, pairwise post-hoc Conover tests between tremor epochs were performed using the “scikit-posthocs” python toolbox. (Terpilowski, 2019) (https://github.com/maximtrp/scikit-posthocs). P-values < 0.05 from post-hoc tests were considered significant.

### Tremor-Neural *θ* Phase Locking Value

To determine whether *θ* (3–8 Hz) in tremor and neural recordings were synchronized, the phase-locking value (PLV) was calculated with tremor and neural *θ* phase per trial (Lachaux et al., 1999). *θ* phase estimates for neural spectral data were calculated by taking the circular/angular mean for narrowband phase estimates between 3–8 Hz at each timepoint (*t*).

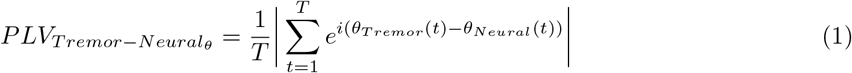

All PLV-related analyses were also calculated with the pairwise phase consistency (PPC) measure to control for differences in number of trials across conditions (Vinck et al., 2010; Aydore et al., 2013).

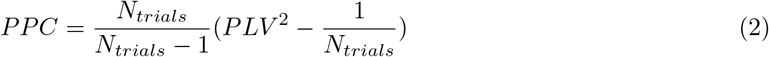

As PLV and PPC results were qualitatively similar, we reported PLV results.

### Tremor-Neural *θ* Phase Slope Index

To understand the lag-lead relationship between tremor (a bandpassed signal) and neural *θ* phase locking, the phase slope index (PSI) was calculated for the *θ* band (3–8 Hz) with 1 Hz frequency resolution (Nolte et al., 2008) using the “spectral_connectivity” python toolbox (https://github.com/Eden-Kramer-Lab/spectral_connectivity, https://doi.org/10.5281/zenodo.4088934).

As the “spectral_connectivity” toolbox uses the multitaper transform for spectral analysis, the number of necessary tapers (*L*) was calculated by first calculating the time-half-bandwidth product (*TW*) using the desired frequency resolution (Δ*f*, 1 Hz for parity with wavelet spectral analyses) and the time window of the entire trial (*N*, 4 seconds) (Prerau et al., 2016).

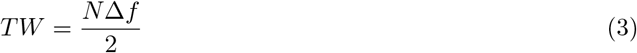

We subsequently used *TW* to calculate the number of tapers (*L*) using the floor function.

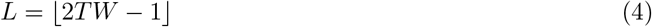

With our parameters, 3 Slepian tapers were used for whole-trial single-window PSI estimates.

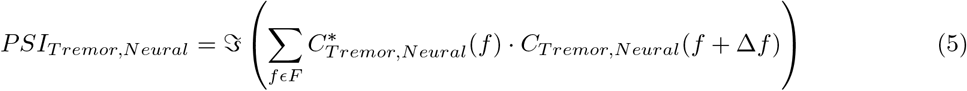

PSI was then estimated from the imaginary 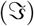 component of the complex coherency (*C*) between tremor and neural *θ*, where the complex coherency was calculated from the cross-spectral density matrix (S) between the two signals.

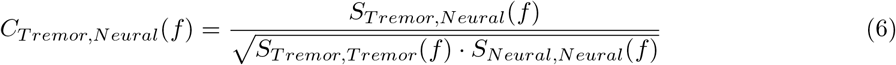

Phase offsets between 1 Hz frequency bands (Δ*f*) within *θ*(*F*) were used to calculate the phase slope. Because of our short-timescale windowed application of PSI, we did not normalize values of PSI by their standard deviation (Young et al., 2017). To determine if tremor or neural recordings exhibited directional *θ* influence, the empirical PSI was compared to a null distribution of 1000 PSI values generated from shuffling one signal’s timeseries across trials. P-values were calculated empirically from the resulting distribution and corrected for multiple comparisons with the Benjamini-Hochberg method at *q* = 0.05.

### Tremor Epoch Interregional Phase Locking Value

To compare time-varying phase synchrony across structures, the phase-locking value (PLV) was calculated across each structure pair (*j, k*) per 1 Hz frequency band (*f*) from 1-100 Hz using wavelet-derived spectral data.

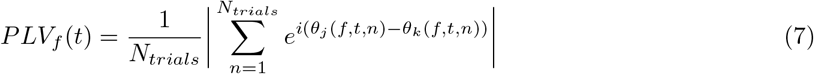

To determine if pairwise PLV differed by tremor epoch type, PLV values within frequencies were averaged across time, and were then compared using the Kruskal-Wallis test. If PLV in a frequency band was found to significantly differ across tremor epochs, pairwise post-hoc Conover tests between tremor epochs were performed (*p* < 0.05 in post-hoc tests were deemed significant).

### Tremor Epoch Interregional Granger Prediction

To understand whether tremor epoch-related dynamic changes in spectral power or synchrony were driven by dynamic directional influences of one structure onto another, nonparametric spectral granger prediction (GP) was calculated between each structure pair using the “spectral_connectivity” python toolbox. Specifically, frequency information (1 Hz frequency resolution) for each structure-timeseries pair were calculated using a single 4000 ms multitaper window (3 tapers). From there, a frequency-based estimation of information flow between structures was calculated using a cross-density spectral matrix (Dhamala et al., 2008). Subsequently, frequency-specific (*f*) GP (i.e. the log-ratio of total frequency power over non-predicted frequency power) was calculated between structure pairs (*j, k*) for each epoch type using the cross-spectral density matrix (*S*), the spectral transfer matrix (*H*), and the noise covariance matrix (Σ).

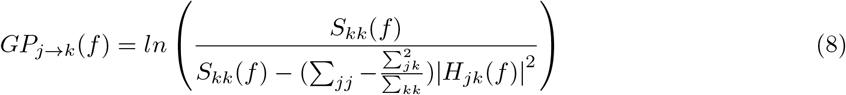

To determine if one structure exhibited frequency-specific granger prediction on another, the empirical GP was compared to a null distribution of 1000 GP values generated from shuffling one structure’s time-series across trials. P-values for each frequency were calculated empirically from the resulting distribution and corrected for multiple comparisons with the Benjamini-Hochberg method at *q* = 0.05.

To understand how GP varied as a function of time, frequency information for each structure-timeseries pair were calculated in 2000 ms windows with 100 ms overlap using the multitaper transform for each event trial. To maintain the same number of tapers (3 tapers) between static and dynamic GP analyses, frequency resolution was increased to 2 Hz for dynamic GP calculation. To determine if one structure exhibited time-varying directional influence on another, the empirical GP was compared to a null distribution of 1000 GP values generated from shuffling one structure’s timeseries across trials. P-values for each time and frequency point were calculated empirically from the resulting distribution and corrected for multiple comparisons with the Benjamini-Hochberg method at *q* = 0.05. Resulting significant time-frequency clusters were additionally filtered by only considering clusters whose area was greater than the 95th percentile of all BH-corrected significant clusters.

### Tremor Epoch Interregional Phase Slope Index

In order to calculate *θ* directed connectivity across structures, the phase slope index (PSI) was used for the *θ* band (3–8 Hz) with 1 Hz frequency resolution across structures. Frequency information (1 Hz frequency resolution) for each structure-timeseries pair were calculated in a single 4000 ms window using the multitaper transform (3 tapers) for each event trial. To determine if one structure exhibited PSI influence on another, the empirical PSI was compared to a null distribution of 1000 PSI values generated from shuffling one structure’s timeseries across trials. P-values were calculated empirically from the resulting distribution and corrected for multiple comparisons with the Benjamini-Hochberg method at *q* = 0.05.

In order to calculate time-varying PSI between broad frequency bands, PSI was calculated using a 2000 ms window sliding by 100 ms (3 tapers with 2 Hz frequency resolution). A bootstrap was then performed, and empirical p-values for each time point were corrected for multiple comparisons with the Benjamini-Hochberg method at *q* = 0.05.

### Statistical Analysis

Data in text are represented as mean ± standard deviation. All statistical tests, unless otherwise specified, were carried out in the “scipy” python environment. P-values were controlled for multiple comparisons by using the Benjamini-Hochberg procedure at *q* = 0.05 (Benjamini and Hochberg, 1995).

### Data and Code Accessibility

The datasets supporting the current study have not been deposited in a public repository because they contain patient information but are available along with analysis code upon request.

## RESULTS

### Intraoperative behavioral and neural data acquisition

Ten patients with PD undergoing DBS implantation and 14 age-matched control subjects (see *Methods*) performed a simple visual-motor task where they followed an onscreen target using a joystick-controlled cursor with their dominant hand **(Figure 1A)**. Each patient performed 1–4 sessions of this target-tracking task during the procedure for a total of 27 sessions, while control subjects performed 1 session each for at total of 14 sessions. Tremor amplitude and cursor speed were quantified continuously from the x- and y-joystick traces. The resulting PD and control tremor and speed distributions were distinct (tremor: *p* = 2.15 × 10^−154^, speed: *p* = 3.44 × 10^−61^, Mann-Whitney U-test) **(Figure 1B-C)**. The partial overlap of the PD and control tremor distributions (indicative of periods without tremor in PD patients), along with the long right tail of the PD distribution, gave us a large dynamic range of tremor to analyze with respect to neural data. The dominant tremor frequency across patients was 4.48 ± 0.57 Hz. While tremor amplitude correlated with the resting tremor UPDRS sub-score (Spearman *ρ* = 0.92, *p* < 0.001, bootstrap test), it did not with the postural tremor sub-score (*ρ* = 0.54, *p* = 0.065, bootstrap test). Based on the distinct tremor frequency peak and its correlation with clinical measures of resting tremor, we interpreted our task-derived tremor as reflective of resting tremor (Dirkx et al., 2018).

Across the 10 patients with PD, we obtained 81 microelectrode recordings within the STN (peak recording density: MNI *x* = +13, *y* = +11, *z* = −5; **Figure 1D**) as well as 72 ECoG recordings from cortex, including premotor cortex (PMC, *n* = 27 recordings), motor cortex (MC, *n* = 16), somatosensory cortex (SC, *n* = 15), and posterior parietal cortex (PPC, *n* = 14) **(Figure 1E)**. All recordings were contralateral to the hand used to perform the task.

### Tremor is a neurophysiologically distinct motor feature of Parkinson’s disease

To understand the relationship of broadband neural activity to tremor expression, we examined the correlation between tremor amplitude and spectral power in neurophysiological recordings. Sorting session-wide spectral data by tremor epochs (rather than according to time) revealed informative band-specific patterns of activity **(Figure 2A)**. Specifically, across cortical structures, spectral power in narrow (1 Hz bandwidth) bands within the *β* range was found to negatively correlate with tremor amplitude (*p* <= 0.008, Spearman *ρ*) **(Figure 2B)**. Interestingly, *β* power appeared to drop off fairly quickly with even low levels of tremor becoming evident (SC - power curve fit : *r*^2^ = 0.77, linear fit : *r*^2^ = 0.54). Meanwhile, *θ* power positively correlated with tremor amplitude in PMC (*p* = 0.005) and SC (*p* <= 0.005). Power in all bands except *β_high_* positively correlated with tremor in STN recordings (*p* <= 0.012).

**Figure 2.**
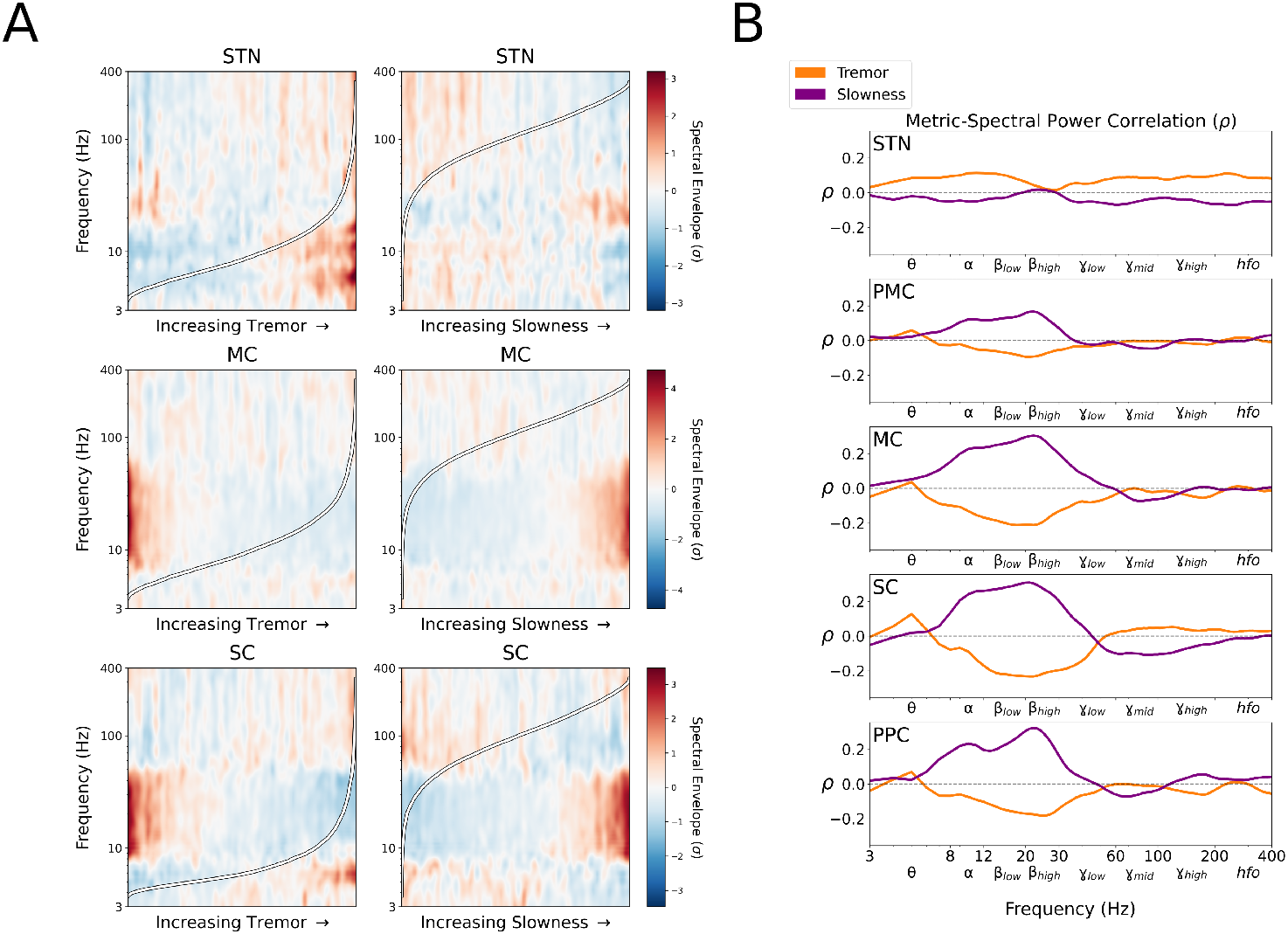
Tremor and slowness exhibit distinct spectral power correlations with intracranial recordings. ***A***, Population-averaged task session spectral power, sorted by each epoch’s tremor amplitude (left) or slowness (right). For ease of visualization, frequency power was Z-scored within frequencies across epochs. ***B***, Average session-wide narrowband (1 Hz) spectral Spearman correlation (ρ) with tremor amplitude and slowness. Note that while *β* frequencies exhibited an opposing relationship with tremor and slowness, *θ* frequencies exhibited a distinct positive correlation with tremor. STN - subthalamic nucleus, PMC - premotor cortex; MC - motor cortex; SC - somatosensory cortex; PPC - parietal cortex.

To compare tremor-related neural activity with a distinct PD motor feature (specifically bradykinesia), neural data were also analyzed with respect to movement “slowness” during the same target-tracking task. Note that PD subjects appeared to lack a higher mode of movement velocity that was clearly present in control subjects, reflecting an inability to move the cursor consistently as quickly as the target **(Figure 1C)**. We calculated a min-max normalized measure of inverse cursor speed (0=highest speed, 1=lowest speed) to capture this effect as a positive pathological sign, parallel to the sign of tremor. In contrast to tremor, we observed positive correlations between slowness and *α*/*β* (8–30 Hz) power in all cortical structures (*p* <= 0.001) **(Figure 2B)**. However, *θ* did not show a significant correlation with slowness in any structure (*p* > 0.05). Thus, *θ* appeared to relate specifically to tremor, whereas the relationship to *β* activity was generally reversed between these PD-related motor manifestations. So while there was broadly the appearance of a symmetric opposition between tremor and slowness in terms of their correlations with neural activity across frequencies **(Figure 2B)**, this difference in the *θ* frequency relationship, as well as perhaps a consistent difference in γ_mid_ (in which the correlation with tremor was typically close to 0 but the correlation with slowness was typically greater in magnitude and negative in direction), suggest these motor features are not simply opposite ends of a single spectrum but rather have distinct fingerprints in neural activity.

### Subthalamic *θ* preceded tremor at onset

Because lower frequency oscillations, particularly *θ*, were most consistently and strongly positively associated with tremor across structures, and because they encompassed the range of observed tremor frequencies from a behavioral perspective (4-6 Hz), we next turned our attention to understanding the relationship of *θ* band activity within each structure to tremor-defined epochs. Using a control vs. PD subject ROC-derived tremor threshold (see *Methods*), behavioral and spectral data were organized into 4 second epochs and categorized as: no tremor epochs (575 epochs, 2300 sec), tremor onset epochs (406 epochs, 1624 sec), and sustained tremor epochs (171 epochs, 684 sec) **(Figure 3A)**. STN *θ* power was indeed significantly elevated during tremor onset and sustained tremor, relative to no tremor (1.07–2.49 fold increase, *p* <= 0.011, Kruskal-Wallis test, *p* < 0.05, Conover test for post-hoc comparisons) **(Figure 3B)**. Likewise, phase synchrony (measured as phase locking value, or PLV) between STN *θ* and tremor was increased during tremor onset and sustained tremor (*p* = 7.20 × 10 ^−39^, Kruskal-Wallis test, *p* < 0.05, Conover test) **(Figure 4A)**.

**Figure 3.**
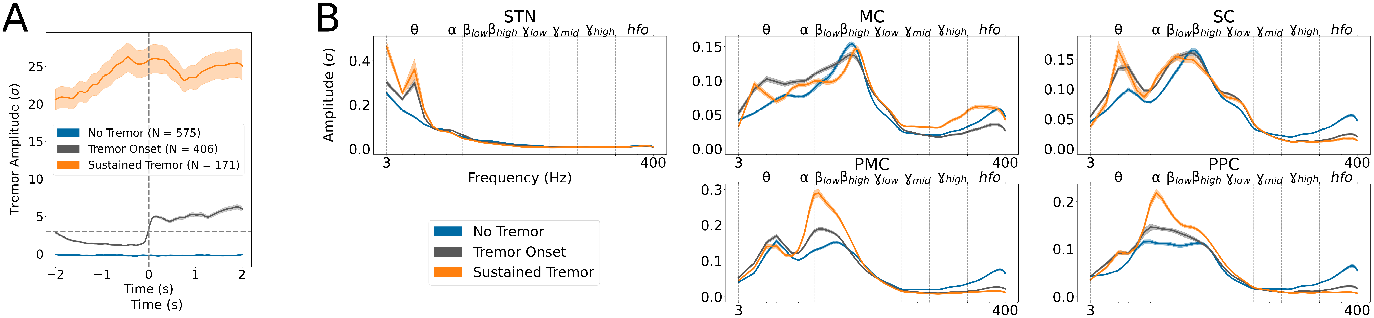
Spectral power during different tremor dynamic states. ***A***, Tremor event design. Based on a population-based tremor ROC threshold, epochs representing different states of tremor dynamics were isolated. For each event type, the average tremor amplitude (±standard error) in patients with PD relative to control subjects is displayed over time. Horizontal dashed line denotes the tremor threshold (3 standard deviations relative to control subjects). Vertical dashed line (*t* = 0) in tremor onset events represents the “trigger” where tremor amplitude crossed the tremor threshold. ***B***, Average spectral power (±standard error) across frequencies for each tremor event type, by recording site. Vertical dashed lines represent frequency band borders. While *θ* oscillations increased in power across STN, MC, and SC, increased tremor was associated with increased *α/β_low_* power in PMC and PPC. STN - subthalamic nucleus, PMC - premotor cortex; MC - motor cortex; SC - somatosensory cortex; PPC - parietal cortex.

**Figure 4.**
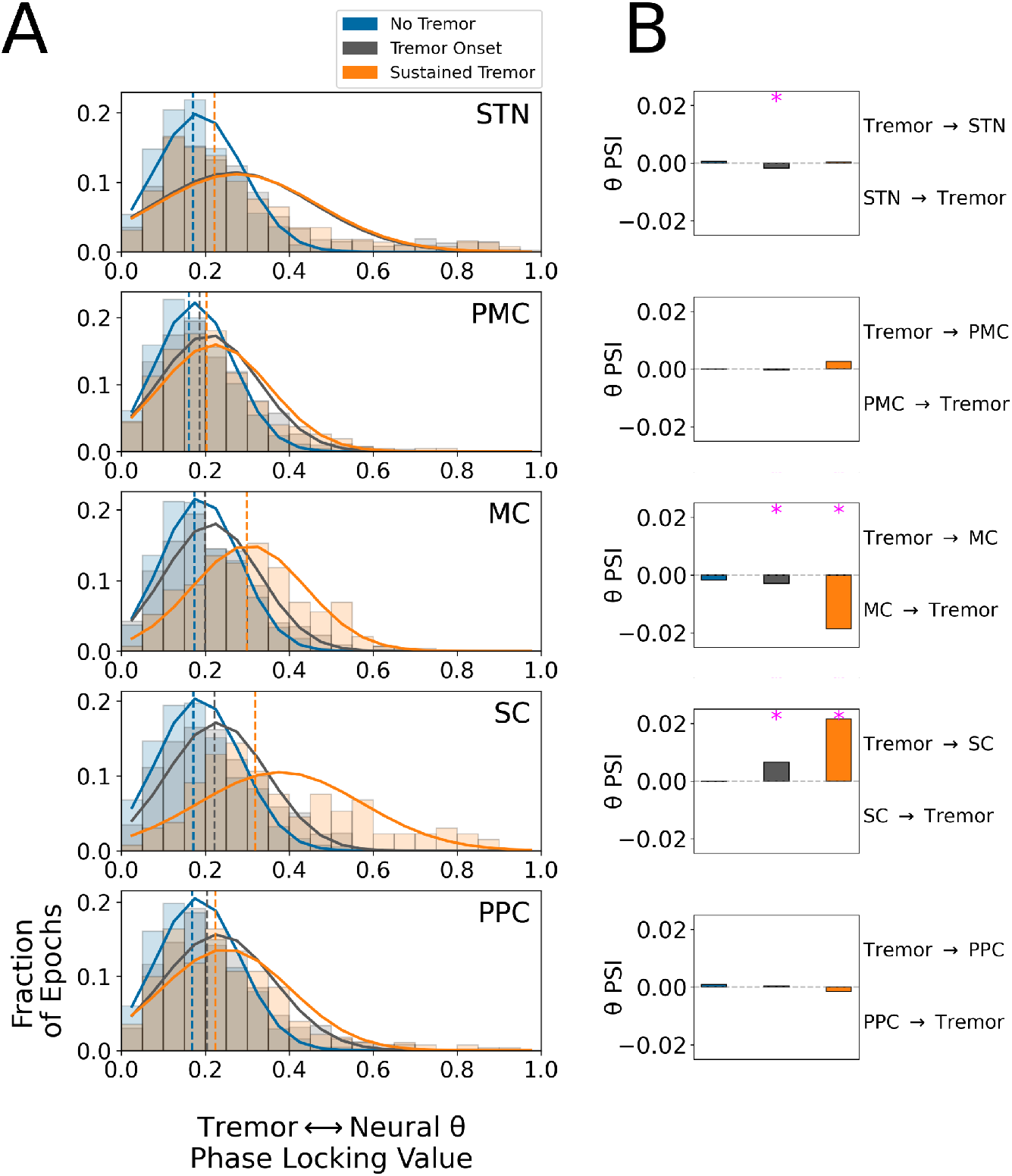
Neural *θ* exhibited structure-specific temporal relationships with tremor. ***A***, Histograms of per-trial phase locking values (PLV) between tremor and neural *θ* by tremor state. Solid lines indicate normal distribution fit to each tremor state PLV histogram, while vertical dashed lines indicate the median of each tremor state PLV histogram. Y-axis indicates proportion of trials within each PLV histogram bin. Note that STN histograms for tremor onset and sustained tremor are highly overlapping. ***B***, Phase slope index (PSI) between tremor and neural *θ* by tremor state. Positive values indicated that tremor phase preceded neural phase, while negative values indicated neural phase preceded tremor. Magenta asterisks indicate significant (*p* < 0.05, bootstrap test) PSI effects. STN - subthalamic nucleus, PMC - premotor cortex; MC - motor cortex; SC - somatosensory cortex; PPC - parietal cortex.

In light of this close relationship between STN *θ* and tremor, we next examined the temporal relationship between STN *θ* and tremor phase. Specifically, we calculated the phase-slope index (PSI) between tremor and STN *θ* phase. Because the PSI considers multiple phase relationships within a range of frequencies, it can succeed in determining the net leading or lagging oscillation in a manner that avoids the circularity problem inherent in methods such as the PLV (Nolte et al., 2008). Here, the PSI revealed STN *θ* led tremor exclusively during tremor onset (*p* = 0.011, bootstrap test) **(Figure 4B)**, consistent with a causal role for the STN in the initiation but not necessarily the maintenance of tremor.

### Somatosensory cortex *θ* consistently followed tremor

Like the STN, SC *θ* power positively correlated with tremor amplitude. Therefore we investigated if this spectral-tremor relationship varied similarly with tremor state. SC *θ* power was indeed significantly elevated during tremor onset and sustained tremor, relative to no tremor (1.08-1.93 fold increase, *p* <= 2.35 × 10^−9^, Kruskal-Wallis test, *p* < 0.05, Conover test) **(Figure 3B)**. SC-tremor *θ* PLV also was increased during tremor onset and sustained tremor (*p* = 4.03 × 10^−37^, Kruskal-Wallis test, *p* < 0.05, Conover test) **(Figure 4A)**.

However, in contrast to the STN, phase-slope analysis of tremor and SC *θ* phase revealed that SC *θ* phase followed tremor phase during both tremor onset and sustained tremor (*p* <= 0.002, bootstrap test) **(Figure 4B)**. Therefore, the strong tremor-related *θ* oscillation seen in SC was reflective rather than causal of tremor.

### Motor cortex *θ* consistently preceded tremor

Unlike the STN and SC, MC *θ* power did not show a clear graded relationship with tremor magnitude **(Figure 2)**. Nonetheless, examining MC *θ* power across tremor states did reveal it was relatively increased when tremor was present, especially during tremor onset (1.20-1.66 fold, *p* <= 0.016, Kruskal-Wallis test, *p* < 0.05, Conover test) **(Figure 3B)**. Furthermore, MC-tremor *θ* PLV increased from no-tremor to tremor-onset to sustained-tremor (*p* = 9.55 × 10^−37^, Kruskal-Wallis test, *p* < 0.05, Conover test) **(Figure 4A)**. Interestingly, examining the PSI for MC *θ* and tremor revealed that MC *θ* led tremor during both tremor onset and sustained tremor (*p* <= 0.014, bootstrap test) **(Figure 4B)**. Thus, in contrast to SC, MC *θ* preceded tremor output.

### Tremor-related *θ* transitioned from STN to cortex during tremor onset

Because both STN and MC *θ* power were elevated during tremor onset, and STN and MC *θ* phase led tremor phase during tremor onset, we investigated the dynamics of STN-MC coupling during the dynamics of tremor initiation. Static phase slope analysis of STN and MC revealed that STN *θ* led MC *θ* during tremor onset (*p* < 0.001, bootstrap test) **(Figure 5A)**. To understand if this phase relationship was time-locked to increasing tremor, we calculated STN-MC *θ* PSI as a function of time within the tremor onset window. Within this epoch, STN *θ* preceded MC *θ* most consistently about 0.5 seconds after tremor detection (*t* = 0) to the end of the tremor onset epoch (*t* = 0.5–1.0 seconds; *p* < 0.05, bootstrap test) **(Figure 5B)**. At no point in this window did MC *θ* appear to precede STN *θ*.

**Figure 5.**
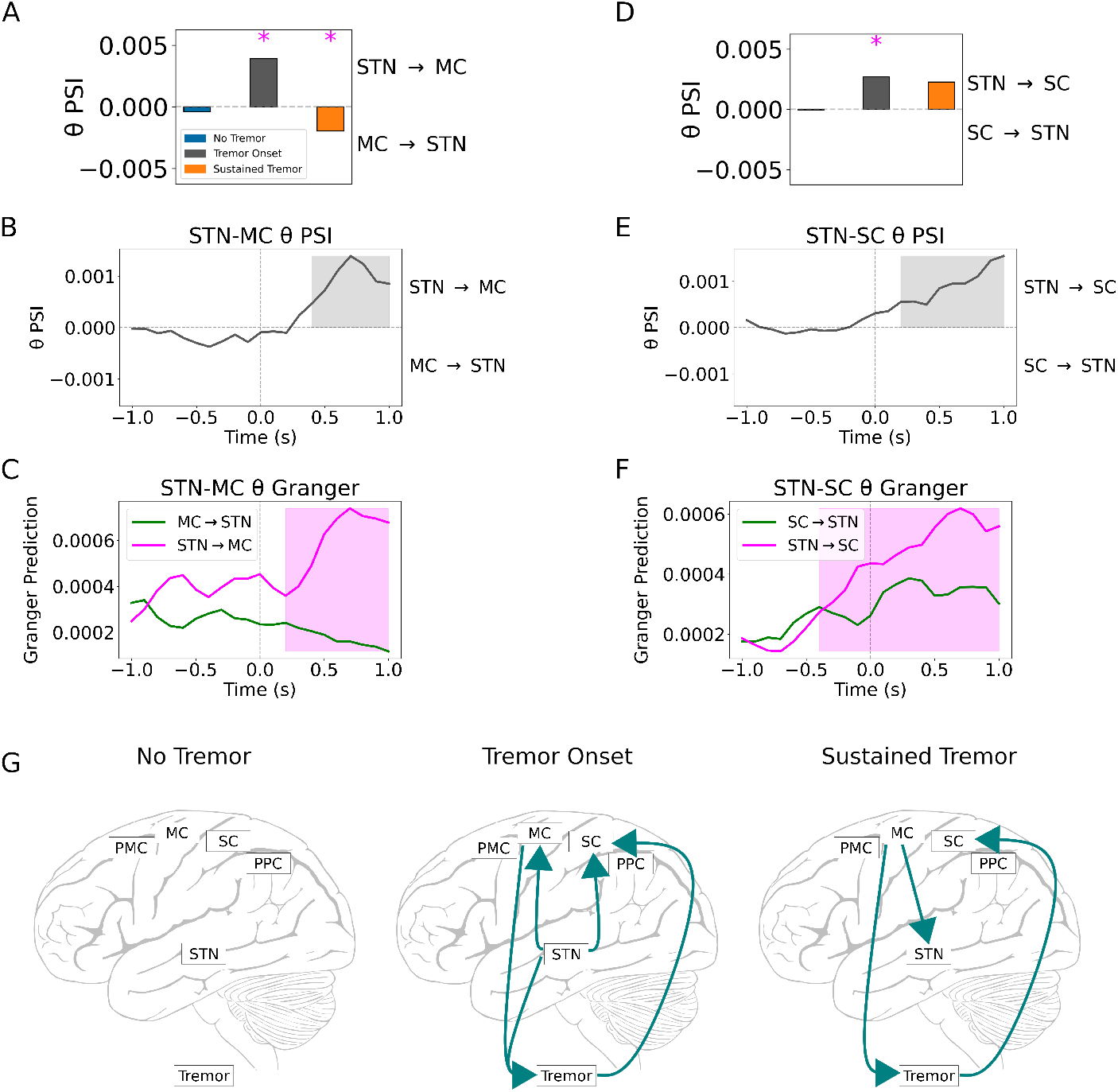
Tremor initiation was driven by the subthalamic nucleus. ***A***, Static phase slope index (PSI) between STN and MC recordings during tremor states. Magenta asterisks indicate significant (*p* < 0.05, bootstrap test) PSI effects. ***B***, Dynamic PSI between STN and MC *θ* during tremor onset. Highlighted regions indicate significant PSI (*p* < 0.05, bootstrap test). Vertical dashed line (*t* = 0) indicates tremor onset trigger. ***C***, Directed granger prediction (GP) between STN and MC *θ* during tremor onset. Vertical dashed line (*t* = 0) indicates tremor onset trigger. Highlighted regions indicate significant granger prediction (*p* < 0.001, bootstrap test). ***D***, Static PSI between STN and SC recordings during tremor states. Magenta asterisks indicate significant (*p* < 0.05, bootstrap test) PSI effects. ***E***, Dynamic PSI between STN and SC *θ* during tremor onset. Highlighted regions indicate significant PSI (*p* < 0.05, bootstrap test). Vertical dashed line (*t* = 0) indicates tremor onset trigger. ***F***, Directed GP between STN and SC *θ* during tremor onset. Vertical dashed line (*t* = 0) indicates tremor onset trigger. Highlighted regions indicate significant granger prediction (*p* < 0.001, bootstrap test). ***G***, Summary of *θ* PSI results. Solid lines represent directed functional connectivity between neural regions and tremor. STN - subthalamic nucleus; PMC - premotor cortex; MC - motor cortex; SC - somatosensory cortex; PPC - parietal cortex.

We also investigated whether STN *θ* and MC *θ* power influenced each other by calculating time-varying nonparametric spectral granger prediction (GP) (see *Methods*). Briefly, a nonzero GP at a particular frequency indicated that spectral power in one structure was predictive of spectral power in another. Unlike the PSI, GP allows the disentangling of asymmetric, bidirectional influences across two signals (Dhamala et al., 2008). As with PSI, STN *θ* power predicted MC *θ* power from 200 ms after the tremor onset trigger to the end of epoch (*t* = 0.2–1.0 seconds; *p* < 0.05, bootstrap test) **(Figure 5C)**. Again, MC *θ* did not predict STN *θ* at any point in the epoch. Together, these results converged to suggest STN *θ* drove MC *θ* during tremor onset.

Once tremor was established however, the *θ* phase slope relationship flipped, with MC *θ* phase preceding STN *θ* phase **(Figure 5A)**, revealing a dynamic transition with increasing tremor. Taken together with the loss of STN *θ* influence over tremor during sustained tremor **(Figure 4B)**, tremor output appeared to become cortically rather than STN driven as tremor became established.

Because the STN and SC both exhibited positive correlations between *θ* power and increasing tremor, we also investigated whether STN/SC dynamics varied during tremor onset. Like MC, static phase slope analysis of STN and SC *θ* revealed that STN *θ* led SC during tremor onset (*p* < 0.001, bootstrap test) **(Figure 5D)**. Dynamic STN-SC PSI additionally revealed that STN *θ* led SC *θ* between 200 ms after the tremor onset trigger to the end of the epoch (*t* = 0.2–1.0 seconds; *p* < 0.001, bootstrap test) **(Figure 5E)**. Simultaneously, STN *θ* power predicted SC *θ* power from 400 ms before the tremor onset trigger to end of the tremor onset epoch (*t* = −0.4−+1.0 seconds; *p* < 0.001, bootstrap test) **(Figure 5F)**. During sustained tremor epochs however, the *θ* phase slope relationship between STN and SC became ambiguous (*p* = 0.091, bootstrap test), again representing a loss of STN influence over cortical *θ* activity **(Figure 5D)**. Altogether, although the STN drove both tremor and cortical *θ* as tremor emerged, the transition to sustained tremor was accompanied by a decoupling of the STN from cortex in the *θ* band **(Figure 5G)**.

### Motor cortex decoupled from posterior cortices with increasing tremor

As STN-MC *θ* phase influence flipped from tremor onset to sustained tremor, we investigated whether the functional connectivity of MC extended to other cortical regions with increasing tremor. To understand if tremor-mediated cortico-cortical interactions occurred in frequency bands other than *θ*, we calculated both nondirected (PLV) and directed (GP) functional connectivity between the MC and other cortical regions across the 3-100 Hz spectrum. While MC-SC PLV was broadly modulated by tremor state (*p* <= 1.81 × 10^−66^, Kruskal-Wallis test), it specifically decreased across all bands except *γ_mid_* during tremor onset (PLV, 1.20-7.72 fold decrease, *p* < 0.05, Conover test) **(Figure 6A)**. To identify whether synchrony detected by the PLV was driven by one structure in the pair, broad-spectrum GP was calculated. However, no consistent band-wide differences in MC-SC GP were found across tremor states (*p* > 0.05, bootstrap) **(Figure 6B)**.

**Figure 6.**
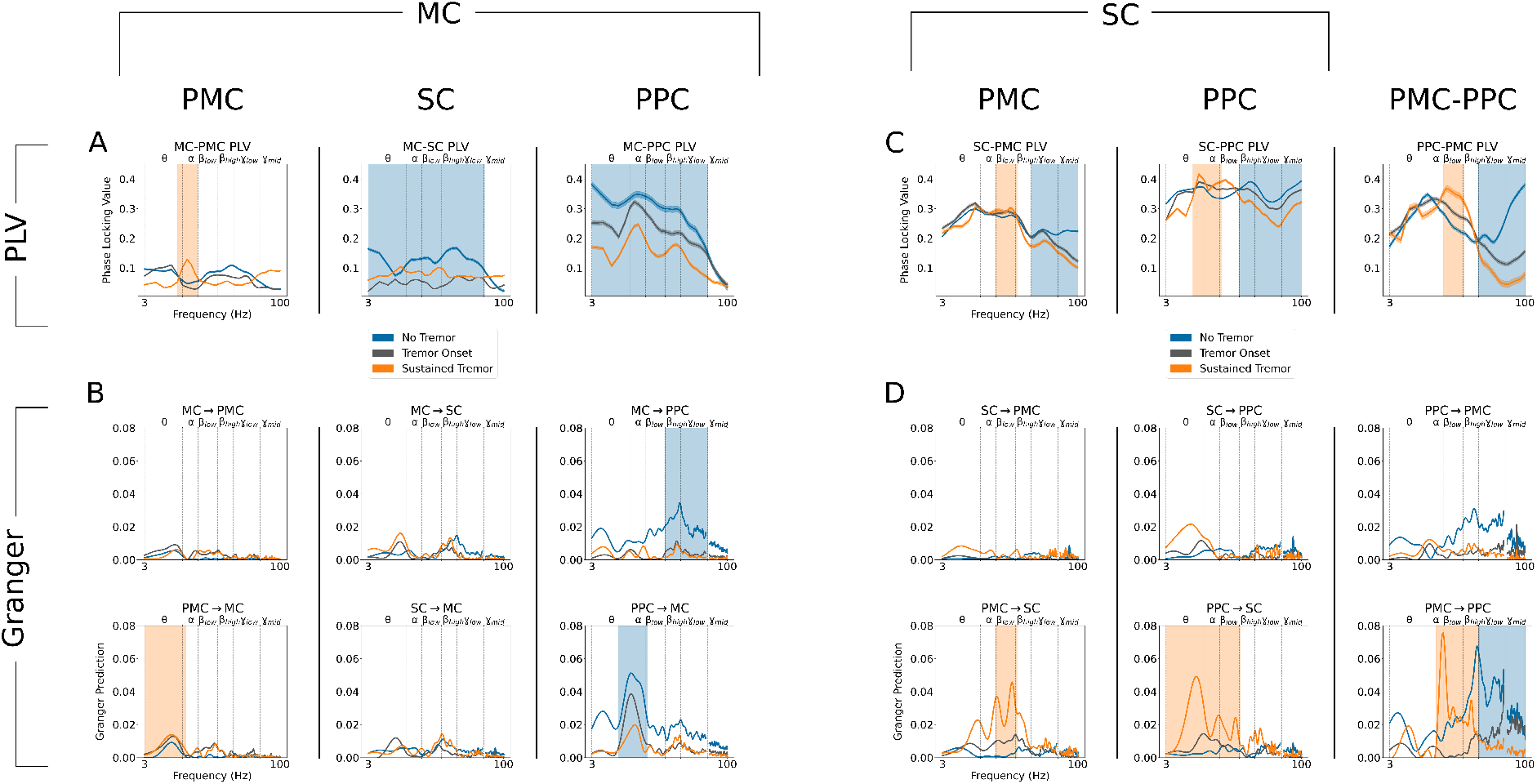
During sustained tremor, gamma coupling between premotor/motor and somatosensory/parietal cortices decreased. ***A***, Phase locking value (PLV) between MC and other cortical regions. Lines *±* shaded borders represent average ± standard error PLV. Highlighted frequency ranges indicate increased (orange) or decreased (blue) PLV with increasing tremor. ***B***, Pairwise granger prediction (GP) between MC and other cortical regions. The title of each subpanel indicates the directionality of the structure pair GP. Highlighted frequency ranges indicate increased (orange) or decreased (blue) GP with increasing tremor. For ease of visualization, curves were lowpass filtered and frequencies within 58–62 Hz were masked. Note that MC broad-spectrum coupling with SC and PPC generally decreased with increasing tremor. ***C***, PLV between SC and other cortical regions. Lines *±* shaded borders represent average ± standard error PLV. Highlighted frequency ranges indicate increased (orange) or decreased (blue) PLV with increasing tremor. ***D***, Pairwise GP between SC and other cortical regions. Title of each subpanel indicates the directionality of the structure pair GP. Highlighted frequency ranges indicate increased (orange) or decreased (blue) GP with increasing tremor. For ease of visualization, curves were lowpass filtered and frequencies within 58–62 Hz were masked. Note that tremor generally shifted the frequency of coupling between SC, PPC, and PMC from *γ* to *α/β_low_* with increasing tremor. Vertical dashed lines represent frequency band borders. STN - subthalamic nucleus; PMC - premotor cortex; MC - motor cortex; SC - somatosensory cortex; PPC - parietal cortex.

MC-PPC PLV similarly decreased across all frequencies except *γ_mid_* as tremor increased (PLV, 1.07–2.95 fold decrease, *p* <= 1.13 × 10^−20^, Kruskal-Wallis test, *p* < 0.05, Conover test) **(Figure 6A)**. Despite this drop, the MC-PPC PLV spectrum revealed coupling peaks in *α* and *β_high_*/*γ_low_* (20-60 Hz) frequencies across all tremor states. However, each peak appeared as a directed channel of communication across MC and PPC, particularly in the absence of tremor. Granger analysis revealed that while PPC *α* predicted MC *α* regardless of tremor state, (GP, 0.94-5.24 fold difference, *p* < 0.001, bootstrap test in all tremor states), its absolute prediction was 3.25 fold smaller in sustained tremor vs. no tremor **(Figure 6B)**. In contrast, MC *β_high_*/*γ_low_* predicted PPC *β_high_*/*γ_low_* exclusively in the absence of tremor (GP, 0.93-1.93 fold difference, *p* < 0.001, bootstrap test).

In sum, MC became less coupled with posterior cortical regions (SC, PPC) with increasing tremor, while MC became increasingly coupled with PMC. Specifically, MC-PMC PLV increased within *θ*/*α* (6-10 Hz) specifically during sustained tremor (PLV, 1.54-3.67 fold increase, *p* <= 1.80 × 10^−152^, Kruskal-Wallis test, *p* < 0.05, Conover test) **(Figure 6A)**. While not entirely within the same frequency range, PMC *θ* appeared to predict MC *θ* during sustained tremor (GP, 1.86-7.28 fold increase, *p* < 0.001, bootstrap test) **(Figure 6B)**.

### Premotor cortex coupled with posterior cortices during tremor

Because SC decoupled from the STN during sustained tremor while still reflecting tremor output, we investigated whether SC instead coupled with other cortical regions as tremor increased. SC and PPC exhibited increased *θ*/*α* (6–12 Hz) PLV (PLV, 1.02–1.16 fold increase, *p* <= 4.62 × 10^−39^, Kruskal-Wallis test, *p* < 0.05, Conover test) and decreased *β_high_-γ* (20–100 Hz) PLV (PLV, 1.10–1.38 fold decrease, *p* <= 1.71 × 10^−91^, Kruskal-Wallis test, *p* < 0.05, Conover test) with increasing tremor **(Figure 6C)**. While SC-PPC functional connectivity was relatively symmetric during low-tremor states (no tremor, tremor onset), sustained tremor revealed more directed coupling. Although *β_low_* PLV did not significantly modulate with tremor, *α*/*β_low_* (8–20 Hz) SC–PPC PLV during sustained tremor was driven by PPC (GP, 1.65–20.85 fold difference, *p* < 0.001, bootstrap test) **(Figure 6D)**. Additionally, PPC *θ* predicted SC *θ* during sustained tremor (GP, 1.73–2.80 fold difference, *p* < 0.001, bootstrap test). Thus, SC-PPC functional connectivity shifted to a distinct state during sustained tremor, with PPC predicting lower frequencies (*θ,α, β_low_*) in SC. At the same time, higher frequency (*γ*) coupling between SC and PPC decreased as tremor increased.

SC and PMC interactions also exhibited push-pull changes in functional connectivity, with increased *β_low_* PLV (PLV, 1.04–1.09 fold increase, *p* <= 6.43 × 10^−33^, Kruskal-Wallis test, *p* < 0.05, Conover test) and decreased *γ* PLV (PLV, 1.42–2.19 fold decrease, *p* <= 1.32 × 10 ^−309^, Kruskal-Wallis test, *p* < 0.05, Conover test) with increasing tremor **(Figure 6C)**. Like PPC, increases in lower frequency PLV (*β_low_*) was driven by PMC specifically during sustained tremor (GP, 4.35–12.26 fold difference, *p* < 0.001, bootstrap test) **(Figure 6D)**.

Thus, in contrast to MC, which broadly decoupled from posterior cortical regions, SC became increasingly coupled with and influenced by both posterior (PPC) and anterior (PMC) cortices with increasing tremor. However, this increase in connectivity was specific to *α*/*β_low_* frequencies while *γ* coupling decreased between SC and PMC/PPC.

To follow the spread of tremor-related cortical coupling, we investigated whether PMC and PPC interacted during sustained tremor. Here, we observed an exaggerated version of the same tremor-induced frequency shift (*γ* to *β*) of power and phase synchrony. When analyzing tremor epoch-related spectral power in PMC and PPC in **Figure 3B**, both regions demonstrated tremor-related increases in *α*/*β_low_* power (PPC : 1.09–1.94 fold increase, *p* <= 3.88 × 10^−29^, Kruskal-Wallis test, *p* < 0.05, Conover test) (PMC: 1.07–2.21 fold increase, *p* <= 4.08 × 10^−14^, Kruskal-Wallis test, *p* < 0.05, Conover test). At the same time PMC and PPC exhibited decreases in *γ_mid_*, *γ_high_*, and *hfo* power relative to no tremor (PMC: 1.22–5.42 fold decrease, *p* <= 1.20 × 10^−57^, Kruskal-Wallis test, *p* < 0.05, Conover test) (PPC : 1.67–7.78 fold decrease, *p* <= 0.011, Kruskal-Wallis test, *p* < 0.05, Conover test).

These similar changes in power were mirrored by changes in PMC-PPC PLV synchrony **(Figure 6C)**. PMC PPC *γ_low–mid_* PLV decreased as tremor increased (PLV, 1.06–6.59 fold decrease, *p* <=1.02 × 10^−232^, Kruskal-Wallis test, *p* < 0.05, Conover test), while PMC-PPC *β_low_* PLV increased with tremor (PLV, 1.15-1.53 fold increase, *p* <= 8.34 × 10^−173^, Kruskal-Wallis test, *p* < 0.05, Conover test). Regardless of tremor state, PMC-PPC phase synchrony was driven by PMC onto PPC. When tremor was absent, PMC *γ* predicted PPC *γ* (GP, 0.94-3.31 fold difference, *p* < 0.001, bootstrap test) **(Figure 6D)**. During sustained tremor, PMC *β* power predicted PPC *β* power (GP, 1.76-12.14 fold difference, *p* < 0.001, bootstrap test).

Overall, tremor was associated with a frequency shift (*γ* to *β*) of power and phase synchrony between PMC, PPC, and SC. Specifically, PMC exerted increasing influence over posterior regions (SC, PPC) in lower frequencies (*α, β_low_*) with increasing tremor. However, this increase in lower frequency coupling coincided with decreases in higher frequency coupling (*γ*). In addition, *γ* coupling between MC and PPC decreased with increasing tremor, revealing that sustained tremor is a state of decreased *γ* synchrony across sensorimotor cortex.

## DISCUSSION

Using a naturalistic behavioral task, we were able to characterize tremor dynamics and isolate specific tremor states, particularly tremor onset and maintenance. Across structures we found that *θ* power positively and *β* power negatively correlated with tremor, as has been found in previous reports (Hirschmann et al., 2013; Qasim et al., 2016; Asch et al., 2020). However, our study is the first to dissect electrophysiological correlates of tremor onset and sustained tremor. During the emergence of tremor, not only did STN and motor cortical *θ* power increase, but STN and motor cortical *θ* phase preceded the phase of tremor. Moreover, STN *θ* activity drove motor cortical *θ* during tremor onset, suggesting a direct role of the STN in initiating tremor output.

Once tremor emerged however, motor cortex appeared to sustain tremor. At the same time, motor cortex became less coupled with somatosensory and parietal cortices, despite the presence of prominent somatosensory cortex *θ* power which closely followed tremor. Instead, premotor cortex synchronized via *β_low_* frequencies with posterior cortices (somatosensory, parietal) at the expense of *γ* frequency synchronization observed in the absence of tremor. This *β_low_* synchrony was notably asymmetric across these structures, with premotor cortex exerting influence over posterior cortices.

Taken together, although tremor amplitude corresponded to global changes in *θ* and *β* power, the relationship between these frequency bands to tremor output was highly structure-specific. While STN-motor cortical interactions appeared to initiate tremor, premotor cortex-driven network effects may help sustain tremor. This STN-mediated dynamic reorganization of cortical connectivity is consistent with both the “dimmer switch” model and the “intrinsic” and “extrinsic” cortical loops of Parkinson’s tremor (Helmich et al., 2011; Volkmann et al., 1996)**(Figure 7)**. Like the GPi, we revealed that the STN acted as a “switch” to mediate the onset of tremor by influencing motor cortex (Dirkx et al., 2016). While these STN-motor cortical interactions formed the “intrinsic” loop of tremor output, we expanded this model to reveal that shifts from *γ* to *β* synchrony across premotor-parietal cortices potentially acted as the “extrinsic” loop to stabilize the tremor state.

**Figure 7.**
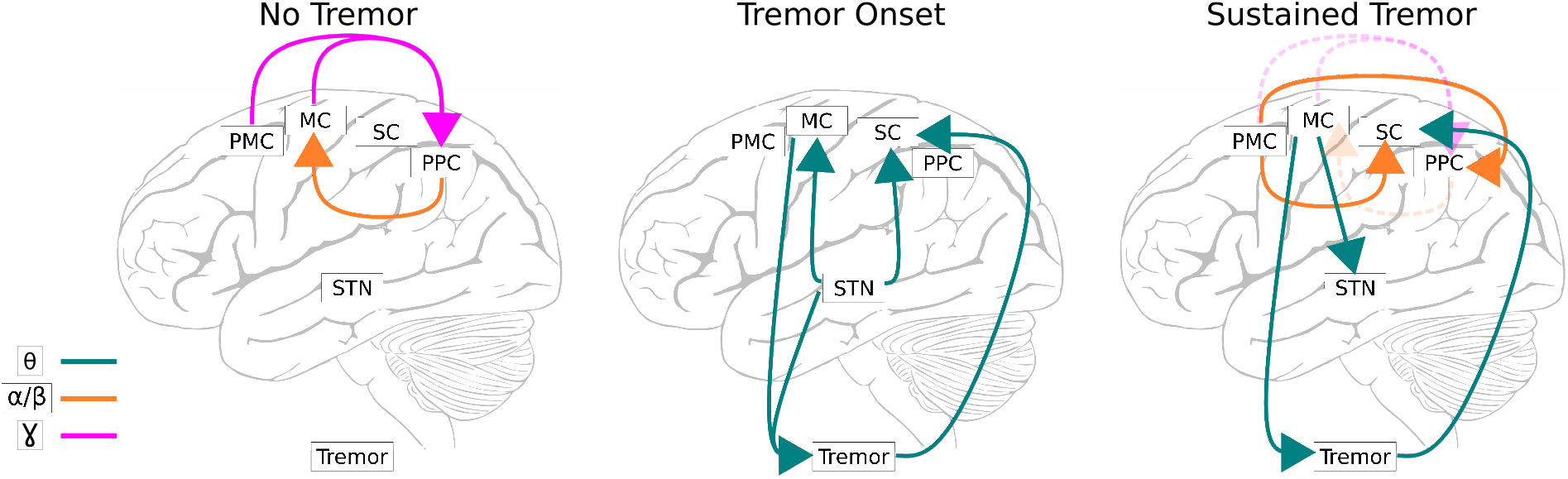
Synthetic model of subcortical-cortical interactions during tremor. Solid lines represent directed functional connectivity between neural regions and tremor. Dashed lines during sustained tremor represent interactions from the no tremor state that are no longer present. STN - subthalamic nucleus; PMC - premotor cortex; MC - motor cortex; SC - somatosensory cortex; PPC - parietal cortex.

### Tremor onset was mediated by subthalamic *θ* driving motor cortex

STN *θ* amplitude positively correlated with tremor amplitude regardless of tremor dynamic states. While the phase of STN *θ* consistently preceded tremor phase during tremor onset, it did not during sustained tremor. However, STN *θ* activity was still significantly phase-locked to tremor during sustained tremor. This mixed relationship to tremor may reflect several roles of STN: interconnections with GPi contribute to tremor initiation, while disynaptic connections with cerebellum may influence ongoing monitoring of tremor output (Helmich et al., 2011; Bostan et al., 2010).

Regardless, STN *θ* drove motor cortex activity during tremor onset. While tremor has previously been found to decrease *β* coherence between STN and motor cortex (Qasim et al., 2016) while increasing *θ* coherence (Hirschmann et al., 2013), we demonstrated directed *θ* phase interactions from STN to motor cortex specifically during tremor onset. While a previous case study of tremor onset displayed local STN and cortical *α*/*β* power changes with tremor onset (Hirschmann et al., 2019), we show here that STN and motor cortical *θ* activity are directionally linked. We also demonstrated that during sustained tremor, the STN-motor cortex *θ* phase slope relationship reversed, suggesting the *θ* influence over sustained tremor shifted source from STN to cortex.

### Motor cortex desynchronized with posterior cortices while sustaining tremor

As tremor progressed, motor cortex *θ* increasingly drove tremor. While previous studies have correlated motor cortical activity to tremor (Helmich et al., 2011; Timmermann et al., 2003), this is the first study to our knowledge that has demonstrated a directed relationship between ECoG recordings and tremor. Although motor cortex was synchronized to tremor, motor cortex appeared to desynchronize with other cortical structures with the exception of premotor cortex, as has been found previously (Timmermann et al., 2003; Qasim et al., 2016). While other studies have found that motor cortex increased its synchrony with premotor and parietal cortices during tremor (Hirschmann et al., 2013), this was calculated only at tremor and double-tremor frequencies.

### Tremor reorganized premotor and parietal cortical coupling

Although premotor and parietal cortices did not exhibit a direct *θ* relationship to tremor, changes in tremor initiated a frequency shift in premotor-parietal coupling dynamics. In the absence of tremor, these regions were functionally coupled at higher frequencies (*β_high_, γ_low–mid_*). fMRI studies in patients with PD have found that these regions exhibit overactive BOLD activity during self-initiated sequential hand movements (Samuel et al., 1997), which is hypothesized to compensate for decreased BOLD activity in fronto-striatal circuits in the dopamine depleted state (Wu et al., 2011). Furthermore, cortical *γ* frequency power and synchrony are associated specifically with voluntary movement (Crone et al., 1998;Miller et al., 2007). In our study, this bidirectional premotor-parietal *γ* activity may have reflected task monitoring and spatial tracking (motor output) using sensory information.

During sustained tremor however, parietal and premotor cortices both exhibited increases in *β_low_* power. This *β_low_* activity was also functionally coupled, with premotor driving parietal cortex. Elevated *β_low_* oscillations have been observed in premotor cortex recordings in MPTP non-human primates with predominantly akinetic/rigid symptoms (Wang et al., 2017). While not observed in our study, increased premotor *β_high_* influence over the STN has also been found to correlate with akinetic/rigid symptoms (Sharott et al., 2018). Premotor *β_low_* oscillations may function here in a similar anti-kinetic fashion with other cortical structures during tremor.

In any case, with increasing tremor premotor-parietal *γ* activity diminished while premotor *β_low_* activity drove parietal activity. These frequency shifts may be best understood in the framework of communication-through-coherence theory (Fries, 2015). Specifically, while symmetric or bottom-up *γ* oscillations permit effective and precise transmission of motor-related information across structures, lower-frequency oscillations such as *α*/*β* act as top-down feedback. Here, task-related *γ* synchrony observed across sensorimotor cortex decreased with tremor. In contrast, lower-frequency oscillations such as *β_low_* increased in synchrony, perhaps acting as pathological “feedback” restricting further voluntary movement. Thus, voluntary movement which normally acts to suppress tremor is impeded (Naros et al.,2018). As motor cortical-thalamo-cerebellar loops have been found to sustain and modulate tremor amplitude (Dirkx et al., 2016), our results extend this model by showing premotor *α*/*β_low_* activity may suppress voluntary movement, allowing tremor to persist.

### Implications for closed-loop deep brain stimulation

Because of the clinical interest in developing adaptive closed-loop DBS to more precisely treat PD symptoms, various electrophysiological observations have been investigated as potential tremor biomarkers to inform stimulation (Hirschmann et al., 2017; Shah et al., 2018; Yao et al., 2020). While promising, the features used for tremor detection do not take into account the dynamic nature of tremor — namely, the distinct neurophysiological signature of tremor onset. Because of the breadth of STN *β*-frequency oscillation research in PD, initial closed-loop DBS efforts have focused on using *β* oscillations as a proxy for bradykinesia symptoms (Little et al., 2013; Little et al., 2016; Little et al., 2016; Velisar et al., 2019). However, *β*-driven DBS has been shown to worsen tremor in some patients (Pia-Fuentes et al., 2020; He et al., 2020).

Here, we demonstrated that subthalamic *θ* was present whether tremor was emerging or sustained. The addition of STN *θ*-based biomarkers to closed-loop DBS could help treat the separate symptom axis of tremor. Further, we have provided the best evidence to date that cortical ECoG *θ* is a robust marker for tremor. Specifically, we found that motor cortical *θ* was synchronized to STN *θ* during tremor states, and that somatosensory *θ* was a reliable indicator of immediate tremor amplitude.

These results overall argue for a combined subcortical-cortical stimulation/recording paradigm not unlike cortical-thalamic closed-loop DBS for ET (Opri et al., 2020). By combining recordings from the STN and sensorimotor cortex, an algorithm could infer whether tremor was about to emerge (STN and MC *θ*) or was already present (SC *θ*). In particular, somatosensory cortical recordings could allow for continuous monitoring of tremor despite any stimulus artifact or competing oscillations in the STN. Ideally, DBS for a patient with a mixed motor phenotype could be optimized between STN *β* for bradykinesia symptoms and SC *θ* oscillations for tremor.

### Limitations and Conclusions

Because all tremor data were quantified from patients as they were moving their upper limb during our tracking task, our tremor conditions do not reflect a pure “rest” tremor. However, as Parkinsonian tremor can often emerge as patients maintain a posture or perform a task, our approach still captured meaningful aspects of tremor. Due to our PD population receiving mostly STN DBS for clinical reasons, we were unable to assess the role of the GPi and motor thalamus (VIM) neurophysiology to tremor onset and/or maintenance. Nevertheless, our awake behaving intraoperative recordings revealed that the STN and motor cortex work together to initiate tremor, and tremor is in part sustained by premotor-parietal synchrony.

## Acknowledgements

We are grateful for the generous participation of our patients in this study. We thank Kelsea Laubenstein-Parker for technical assistance, Karina Bertsch for administrative support, and Ann Duggan-Winkle for clinical support. We also thank Minkyu Ahn, David Segar, Tina Sankhla, and Daniel Shiebler for helping develop the motor task experiment. This work was supported by an NIH Training Grant (NINDS T32MH020068) to P.M.L., a Doris Duke Clinical Scientist Development Award (#2014101) to W.F.A., an NIH COBRE Award: NIGMS P20 GM103645 (PI: Jerome Sanes) supporting W.F.A., a Neurosurgery Research and Education Foundation (NREF) grant to W.F.A., the Lifespan Norman Prince Neurosciences Institute, and the Brown University Robert J. and Nancy D. Carney Institute for Brain Science. Part of this research was conducted using computational resources and services at the Center for Computation and Visualization at Brown University, with funding provided by an NIH Office of the Director grant S10OD025181. W.F.A. has received proprietary equipment and technical support for unrelated research through the Medtronic external research program.

## REFERENCES

Accolla E, Caputo E, Cogiamanian F, Tamma F, MrakicSposta S, Marceglia S, Egidi M, Rampini P, Locatelli M, Priori A (2007) Gender differences in patients with Parkinson’s disease treated with subthalamic deep brain stimulation. Movement Disorders 22:1150–1156 _eprint: https://onlinelibrary.wiley.com/doi/pdf/10.1002/mds.21520.

Akbar U, Asaad WF (2017) A Comprehensive Approach to Deep Brain Stimulation for Movement Disorders. Rhode Island Medical Journal (2013) 100:30–33.

Albin RL, Young AB, Penney JB (1989) The functional anatomy of basal ganglia disorders. Trends in Neurosciences 12:366–375.

Argall BD, Saad ZS, Beauchamp MS (2006) Simplified intersubject averaging on the cortical surface using SUMA. Human Brain Mapping 27:14–27 _eprint: https://onlinelibrary.wiley.com/doi/pdf/10.1002/hbm.20158.

Asaad WF, Eskandar EN (2008a) Achieving behavioral control with millisecond resolution in a high-level programming environment. Journal of neuroscience methods 173:235–240.

Asaad WF, Eskandar EN (2008b) A flexible software tool for temporally-precise behavioral control in Matlab. Journal of neuroscience methods 174:245–258.

Asaad WF, Santhanam N, McClellan S, Freedman DJ (2013) High-performance execution of psychophysical tasks with complex visual stimuli in MATLAB. Journal of Neurophysiology 109:249–260.

Asch N, Herschman Y, Maoz R, Aurbach-Asch C, Valsky D, Abu-Snineh M, Arkadir D, Linetsky E, Eitan R, Marmor O, Bergman H, Israel Z (2020) Independently together: subthalamic theta and beta opposite roles in predicting Parkinsons tremor. Brain Communications.

Aydore S, Pantazis D, Leahy RM (2013) A note on the phase locking value and its properties. NeuroImage 74:231–244.

Benjamini Y, Hochberg Y (1995) Controlling the False Discovery Rate: A Practical and Powerful Approach to Multiple Testing. Journal of the Royal Statistical Society. Series B (Methodological) 57:289–300.

Bergman H, Wichmann T, Karmon B, DeLong MR (1994) The primate subthalamic nucleus. II. Neuronal activity in the MPTP model of parkinsonism. Journal of Neurophysiology 72:507–520.

Bostan AC, Dum RP, Strick PL (2010) The basal ganglia communicate with the cerebellum. Proceedings of the National Academy of Sciences 107:8452–8456 Publisher: National Academy of Sciences Section: Biological Sciences.

Cagnan H, Little S, Foltynie T, Limousin P, Zrinzo L, Hariz M, Cheeran B, Fitzgerald J, Green AL, Aziz T, Brown P (2014) The nature of tremor circuits in parkinsonian and essential tremor. Brain 137:3223–3234.

Cox RW (1996) AFNI: Software for Analysis and Visualization of Functional Magnetic Resonance Neuroimages. Computers and Biomedical Research 29:162–173.

Crone NE, Miglioretti DL, Gordon B, Lesser RP (1998) Functional mapping of human sensorimotor cortex with electrocorticographic spectral analysis. II. Event-related synchronization in the gamma band. Brain 121:2301–2315.

Dale AM, Fischl B, Sereno MI (1999) Cortical Surface-Based Analysis: I. Segmentation and Surface Reconstruction. NeuroImage 9:179–194.

Desikan RS, Sgonne F, Fischl B, Quinn BT, Dickerson BC, Blacker D, Buckner RL, Dale AM, Maguire RP, Hyman BT, Albert MS, Killiany RJ (2006) An automated labeling system for subdividing the human cerebral cortex on MRI scans into gyral based regions of interest. NeuroImage 31:968–980.

Destrieux C, Fischl B, Dale A, Halgren E (2010) Automatic parcellation of human cortical gyri and sulci using standard anatomical nomenclature. NeuroImage 53:1–15.

Dhamala M, Rangarajan G, Ding M (2008) Analyzing Information Flow in Brain Networks with Nonparametric Granger Causality. NeuroImage 41:354–362.

Dirkx MF, den Ouden HEM, Aarts E, Timmer MHM, Bloem BR, Toni I, Helmich RC (2017) Dopamine controls Parkinsons tremor by inhibiting the cerebellar thalamus. Brain 140:721–734.

Dirkx MF, Ouden Hd, Aarts E, Timmer M, Bloem BR, Toni I, Helmich RC (2016) The Cerebral Network of Parkinson’s Tremor: An Effective Connectivity fMRI Study. Journal of Neuroscience 36:5362–5372 Publisher: Society for Neuroscience Section: Articles.

Dirkx MF, Zach H, Bloem BR, Hallett M, Helmich RC (2018) The nature of postural tremor in Parkinson disease. Neurology 90:e1095–e1103.

Dirkx MF, Zach H, van Nuland A, Bloem BR, Toni I, Helmich RC (2019) Cerebral differences between dopamine-resistant and dopamine-responsive Parkinsons tremor. Brain 142:3144–3157.

Fischl B, Salat DH, Busa E, Albert M, Dieterich M, Haselgrove C, van der Kouwe A, Killiany R, Kennedy D, Klaveness S, Montillo A, Makris N, Rosen B, Dale AM (2002) Whole Brain Segmentation: Automated Labeling of Neuroanatomical Structures in the Human Brain. Neuron 33:341–355.

Fonov V, Evans A, McKinstry R, Almli C, Collins D (2009) Unbiased nonlinear average age-appropriate brain templates from birth to adulthood. NeuroImage 47:S102.

Fries P (2015) Rhythms for Cognition: Communication through Coherence. Neuron 88:220–235.

Gross RE, Krack P, Rodriguez-Oroz MC, Rezai AR, Benabid AL (2006) Electrophysiological mapping for the implantation of deep brain stimulators for Parkinson’s disease and tremor. Movement Disorders: Official Journal of the Movement Disorder Society 21 Suppl 14:S259–283.

Hariz GM, Nakajima T, Limousin P, Foltynie T, Zrinzo L, Jahanshahi M, Hamberg K (2011) Gender distribution of patients with Parkinsons disease treated with subthalamic deep brain stimulation; a review of the 20002009 literature. Parkinsonism & Related Disorders 17:146–149.

Harris CR, Millman KJ, van der Walt SJ, Gommers R, Virtanen P, Cournapeau D, Wieser E, Taylor J, Berg S, Smith NJ, Kern R, Picus M, Hoyer S, van Kerkwijk MH, Brett M, Haldane A, del Ro JF, Wiebe M, Peterson P, Grard-Marchant P, Sheppard K, Reddy T, Weckesser W, Abbasi H, Gohlke C, Oliphant TE (2020) Array programming with NumPy. Nature 585:357–362 Number: 7825 Publisher: Nature Publishing Group.

He S, Mostofi A, Syed E, Torrecillos F, Tinkhauser G, Fischer P, Pogosyan A, Hasegawa H, Li Y, Ashkan K, Pereira E, Brown P, Tan H (2020) Subthalamic beta targeted neurofeedback speeds up movement initiation but increases tremor in Parkinsonian patients. eLife 9:e60979 Publisher: eLife Sciences Publications, Ltd.

Helmich RC (2018) The cerebral basis of Parkinsonian tremor: A network perspective. Movement Disorders 33:219–231.

Helmich RC, Hallett M, Deuschl G, Toni I, Bloem BR (2012) Cerebral causes and consequences of parkinsonian resting tremor: a tale of two circuits? Brain 135:3206–3226.

Helmich RC, Janssen MJR, Oyen WJG, Bloem BR, Toni I (2011) Pallidal dysfunction drives a cerebellothalamic circuit into Parkinson tremor. Annals of Neurology 69:269–281.

Hirschmann J, Schoffelen JM, Schnitzler A, van Gerven MAJ (2017) Parkinsonian rest tremor can be detected accurately based on neuronal oscillations recorded from the subthalamic nucleus. Clinical Neurophysiology 128:2029–2036.

Hirschmann J, Abbasi O, Storzer L, Butz M, Hartmann CJ, Wojtecki L, Schnitzler A (2019) Longitudinal Recordings Reveal Transient Increase of Alpha/Low-Beta Power in the Subthalamic Nucleus Associated With the Onset of Parkinsonian Rest Tremor. Frontiers in Neurology 10 Publisher: Frontiers.

Hirschmann J, Butz M, Hartmann CJ, Hoogenboom N, zkurt TE, Vesper J, Wojtecki L, Schnitzler A (2016) Parkinsonian Rest Tremor Is Associated With Modulations of Subthalamic High-Frequency Oscillations. Movement Disorders 31:1551–1559.

Hirschmann J, Hartmann CJ, Butz M, Hoogenboom N, zkurt TE, Elben S, Vesper J, Wojtecki L, Schnitzler A (2013) A direct relationship between oscillatory subthalamic nucleuscortex coupling and rest tremor in Parkinsons disease. Brain 136:3659–3670.

Jankovic J, McDermott M, Carter J, Gauthier S, Goetz C, Golbe L, Huber S, Koller W, Olanow C, Shoulson I (1990) Variable expression of Parkinson’s disease: a base-line analysis of the DATATOP cohort. The Parkinson Study Group. Neurology 40:1529–1534.

Koller WC (1984) The Diagnosis of Parkinson’s Disease. Archives of Internal Medicine 144:2146–2147 Publisher: American Medical Association.

Koller WC (1986) Pharmacologic Treatment of Parkinsonian Tremor. Archives of Neurology 43:126–127 Publisher: American Medical Association.

Konrad PE, Neimat JS, Yu H, Kao CC, Remple MS, D’Haese PF, Dawant BM (2011) Customized, miniature rapid-prototype stereotactic frames for use in deep brain stimulator surgery: initial clinical methodology and experience from 263 patients from 2002 to 2008. Stereotactic and Functional Neurosurgery 89:34–41.

Lachaux JP, Rodriguez E, Martinerie J, Varela FJ (1999) Measuring phase synchrony in brain signals. Human Brain Mapping 8:194–208.

Lance JW, Schwab RS, Peterson EA (1963) ACTION TREMOR AND THE COGWHEEL PHENOMENON IN PARKINSONS DISEASE. Brain 86:95–110 Publisher: Oxford Academic.

Lauro PM, Lee S, Ahn M, Barborica A, Asaad WF (2018) DBStar: An Open-Source Tool Kit for Imaging Analysis with Patient-Customized Deep Brain Stimulation Platforms. Stereotactic and Functional Neurosurgery.

Lauro PM, Vanegas-Arroyave N, Huang L, Taylor PA, Zaghloul KA, Lungu C, Saad ZS, Horovitz SG (2015) DBSproc: An open source process for DBS electrode localization and tractographic analysis. Human Brain Mapping.

Levy R, Hutchison WD, Lozano AM, Dostrovsky JO (2000) High-frequency Synchronization of Neuronal Activity in the Subthalamic Nucleus of Parkinsonian Patients with Limb Tremor. Journal of Neuroscience 20:7766–7775.

Li X, Morgan PS, Ashburner J, Smith J, Rorden C (2016) The first step for neuroimaging data analysis: DICOM to NIfTI conversion. Journal of Neuroscience Methods 264:47–56.

Little S, Beudel M, Zrinzo L, Foltynie T, Limousin P, Hariz M, Neal S, Cheeran B, Cagnan H, Gratwicke J, Aziz TZ, Pogosyan A, Brown P (2016) Bilateral adaptive deep brain stimulation is effective in Parkinson’s disease. Journal of Neurology, Neurosurgery, and Psychiatry 87:717–721.

Little S, Pogosyan A, Neal S, Zavala B, Zrinzo L, Hariz M, Foltynie T, Limousin P, Ashkan K, FitzGerald J, Green AL, Aziz TZ, Brown P (2013) Adaptive deep brain stimulation in advanced Parkinson disease. Annals of Neurology 74:449–457.

Little S, Tripoliti E, Beudel M, Pogosyan A, Cagnan H, Herz D, Bestmann S, Aziz T, Cheeran B, Zrinzo L, Hariz M, Hyam J, Limousin P, Foltynie T, Brown P (2016) Adaptive deep brain stimulation for Parkinson’s disease demonstrates reduced speech side effects compared to conventional stimulation in the acute setting. Journal of Neurology, Neurosurgery, and Psychiatry 87:1388–1389.

Liu Y, Coon WG, Pesters Ad, Brunner P, Schalk G (2015) The effects of spatial filtering and artifacts on electrocorticographic signals. Journal of Neural Engineering 12:056008 Publisher: IOP Publishing.

Miller KJ, Leuthardt EC, Schalk G, Rao RPN, Anderson NR, Moran DW, Miller JW, Ojemann JG (2007) Spectral Changes in Cortical Surface Potentials during Motor Movement. Journal of Neuroscience 27:2424–2432.

Naros G, Grimm F, Weiss D, Gharabaghi A (2018) Directional communication during movement execution interferes with tremor in Parkinson’s disease. Movement Disorders 33:251–261 _eprint: https://onlinelibrary.wiley.com/doi/pdf/10.1002/mds.27221.

Nolte G, Ziehe A, Nikulin VV, Schlgl A, Krmer N, Brismar T, Mller KR (2008) Robustly Estimating the Flow Direction of Information in Complex Physical Systems. Physical Review Letters 100:234101 Publisher: American Physical Society.

Opri E, Cernera S, Molina R, Eisinger RS, Cagle JN, Almeida L, Denison T, Okun MS, Foote KD, Gunduz A (2020) Chronic embedded cortico-thalamic closed-loop deep brain stimulation for the treatment of essential tremor. Science Translational Medicine 12 Publisher: American Association for the Advancement of Science Section: Research Article.

Pia-Fuentes D, van Dijk JMC, van Zijl JC, Moes HR, van Laar T, Oterdoom DLM, Little S, Brown P, Beudel M (2020) Acute effects of Adaptive Deep Brain Stimulation in Parkinsons disease. Brain Stimulation.

Prerau MJ, Brown RE, Bianchi MT, Ellenbogen JM, Purdon PL (2016) Sleep Neurophysiological Dynamics Through the Lens of Multitaper Spectral Analysis. Physiology 32:60–92 Publisher: American Physiological Society.

Qasim SE, de Hemptinne C, Swann NC, Miocinovic S, Ostrem JL, Starr PA (2016) Electrocorticography reveals beta desynchronization in the basal ganglia-cortical loop during rest tremor in Parkinson’s disease. Neurobiology of Disease 86:177–186.

Raz A, Vaadia E, Bergman H (2000) Firing Patterns and Correlations of Spontaneous Discharge of Pallidal Neurons in the Normal and the Tremulous 1-Methyl-4-Phenyl-1,2,3,6-Tetrahydropyridine Vervet Model of Parkinsonism. Journal of Neuroscience 20:8559–8571 Publisher: Society for Neuroscience Section: ARTICLE.

Reck C, Florin E, Wojtecki L, Krause H, Groiss S, Voges J, Maarouf M, Sturm V, Schnitzler A, Timmermann L (2009) Characterisation of tremor-associated local field potentials in the subthalamic nucleus in Parkinsons disease. European Journal of Neuroscience 29:599–612.

Reck C, Himmel M, Florin E, Maarouf M, Sturm V, Wojtecki L, Schnitzler A, Fink GR, Timmermann L (2010) Coherence analysis of local field potentials in the subthalamic nucleus: differences in parkinsonian rest and postural tremor. European Journal of Neuroscience 32:1202–1214.

Rumalla K, Smith KA, Follett KA, Nazzaro JM, Arnold PM (2018) Rates, causes, risk factors, and outcomes of readmission following deep brain stimulation for movement disorders: Analysis of the U.S. Nationwide Readmissions Database. Clinical Neurology and Neurosurgery 171:129–134.

Saad ZS, Reynolds RC (2012) SUMA. NeuroImage 62:768–773.

Samuel M, Ceballos-Baumann AO, Blin J, Uema T, Boecker H, Passingham RE, Brooks DJ (1997) Evidence for lateral premotor and parietal overactivity in Parkinson’s disease during sequential and bimanual movements. A PET study. Brain 120:963–976 Publisher: Oxford Academic.

Shah SA, Tinkhauser G, Chen CC, Little S, Brown P (2018) Parkinsonian Tremor Detection from Subthalamic Nucleus Local Field Potentials for Closed-Loop Deep Brain Stimulation. Conference proceedings: … Annual International Conference of the IEEE Engineering in Medicine and Biology Society. IEEE Engineering in Medicine and Biology Society. Annual Conference 2018:2320–2324.

Sharott A, Gulberti A, Hamel W, Kppen JA, Mnchau A, Buhmann C, Ptter-Nerger M, Westphal M, Gerloff C, Moll CKE, Engel AK (2018) Spatio-temporal dynamics of cortical drive to human subthalamic nucleus neurons in Parkinson’s disease. Neurobiology of Disease 112:49–62.

Telkes I, Viswanathan A, Jimenez-Shahed J, Abosch A, Ozturk M, Gupte A, Jankovic J, Ince NF (2018) Local field potentials of subthalamic nucleus contain electrophysiological footprints of motor subtypes of Parkinson’s disease. Proceedings of the National Academy of Sciences of the United States of America 115:E8567–E8576.

Terpilowski M (2019) scikit-posthocs: Pairwise multiple comparison tests in Python. Journal of Open Source Software 4:1169.

Timmermann L, Gross J, Dirks M, Volkmann J, Freund HJ, Schnitzler A (2003) The cerebral oscillatory network of parkinsonian resting tremor. Brain 126:199–212.

Trotta MS, Cocjin J, Whitehead E, Damera S, Wittig JH, Saad ZS, Inati SK, Zaghloul KA (2018) Surface based electrode localization and standardized regions of interest for intracranial EEG. Human Brain Mapping 39:709–721.

Velisar A, Syrkin-Nikolau J, Blumenfeld Z, Trager MH, Afzal MF, Prabhakar V, Bronte-Stewart H (2019) Dual threshold neural closed loop deep brain stimulation in Parkinson disease patients. Brain Stimulation.

Vinck M, van Wingerden M, Womelsdorf T, Fries P, Pennartz CMA (2010) The pairwise phase consistency: A bias-free measure of rhythmic neuronal synchronization. NeuroImage 51:112–122.

Virtanen P, Gommers R, Oliphant TE, Haberland M, Reddy T, Cournapeau D, Burovski E, Peterson P, Weckesser W, Bright J, Walt SJvd, Brett M, Wilson J, Millman KJ, Mayorov N, Nelson ARJ, Jones E, Kern R, Larson E, Carey CJ, Polat, Feng Y, Moore EW, VanderPlas J, Laxalde D, Perktold J, Cimrman R, Henriksen I, Quintero EA, Harris CR, Archibald AM, Ribeiro AH, Pedregosa F, Mulbregt Pv (2020) SciPy 1.0: fundamental algorithms for scientific computing in Python. Nature Methods pp. 1–12.

Volkmann J, Joliot M, Mogilner A, Ioannides AA, Lado F, Fazzini E, Ribary U, Llinas R (1996) Central motor loop oscillations in parkinsonian resting tremor revealed magnetoencephalography. Neurology 46:1359–1359.

Wang J, Johnson LA, Jensen AL, Baker KB, Molnar GF, Johnson MD, Vitek JL (2017) Network-wide oscillations in the parkinsonian state: alterations in neuronal activities occur in the premotor cortex in parkinsonian nonhuman primates. Journal of Neurophysiology 117:2242–2249 Publisher: American Physiological Society.

Wong JK, Viswanathan VT, Nozile-Firth KS, Eisinger RS, Leone EL, Desai AM, Foote KD, Ramirez-Zamora A, Okun MS, Wagle Shukla A (2020) STN Versus GPi Ddeep Brain Stimulation for Action and Rest Tremor in Parkinsons Disease. Frontiers in Human Neuroscience 14 Publisher: Frontiers.

Wu T, Wang L, Hallett M, Chen Y, Li K, Chan P (2011) Effective connectivity of brain networks during self-initiated movement in Parkinson’s disease. NeuroImage 55:204–215.

Xiao Y, Beriault S, Pike GB, Collins DL (2012) Multicontrast multiecho FLASH MRI for targeting the subthalamic nucleus. Magnetic Resonance Imaging 30:627–640.

Xiao Y, Fonov V, Briault S, Al Subaie F, Chakravarty MM, Sadikot AF, Pike GB, Collins DL (2015) Multi-contrast unbiased MRI atlas of a Parkinson’s disease population. International Journal of Computer Assisted Radiology and Surgery 10:329–341.

Xiao Y, Fonov V, Chakravarty MM, Beriault S, Al Subaie F, Sadikot A, Pike GB, Bertrand G, Collins DL (2017) A dataset of multi-contrast population-averaged brain MRI atlases of a Parkinson’s disease cohort. Data in Brief 12:370–379.

Yao L, Brown P, Shoaran M (2020) Improved detection of Parkinsonian resting tremor with feature engineering and Kalman filtering. Clinical Neurophysiology 131:274–284.

Young CK, Ruan M, McNaughton N (2017) A Critical Assessment of Directed Connectivity Estimates with Artificially Imposed Causality in the Supramammillary-Septo-Hippocampal Circuit. Frontiers in Systems Neuroscience 11.

Zach H, Dirkx M, Bloem BR, Helmich RC (2015) The Clinical Evaluation of Parkinsons Tremor. Journal of Parkinson’s Disease 5:471–474 Publisher: IOS Press.

